# Heparin MicroIslands to Promote Enhanced Diabetic Wound Healing Outcomes

**DOI:** 10.1101/2020.10.31.363531

**Authors:** Lauren Pruett, Christian Jenkins, Neharika Singh, Katarina Catallo, Donald Griffin

## Abstract

A powerful tool to improve tissue integration with biomaterial scaffolds for the regeneration of damaged tissues is to promote cell migration using chemotactic gradients of growth factors. This approach has been realized by the exogenous delivery of growth factors, which unfortunately also limits the scaffold’s ability to meet each wound’s unique spatial and temporal regenerative needs. To address this limitation, we present a new approach to gradient generation by incorporating heparin microislands, which are spatially isolated heparin-containing microparticles that create chemotactic microgradients through reorganization of endogenous local growth factors. We incorporated heparin microislands within microporous annealed particle (MAP) scaffolds, which allows us to tune their incorporation ratiometrically to create a heterogenous microenvironment. In this manuscript, we demonstrate the ability of heparin microislands to organize uniform growth factors into spontaneous microgradients and control downstream cell migration *in vitro*. Further, we present their ability to significantly improve wound healing outcomes (epidermal regeneration and vascularization) in a diabetic wound model relative to two clinically relevant controls.

## Introduction

Growth factors are key regulators of each stage of wound repair, including cellular migration, proliferation, angiogenesis, and extracellular matrix remodeling^1^. Their potential for accelerating wound healing has led to a long history of research efforts focused on biomaterial scaffold-mediated delivery of a range of growth factors (e.g. PDGF, VEGF, EGF, FGF)^2–4^. Importantly, despite FDA approval, growth factor delivery has been limited in clinical translation by safety concerns and cost-effectiveness^2,3,5,6^.

In uninjured tissues, growth factors are an essential component of the instructional microenvironment and their spatial organization within that microenvironment is shaped, in part, by their affinity interactions with the structural components of the extracellular matrix (ECM)^1,2^. Specifically, the ECM can regulate growth factor movement and create chemotactic gradients for spatio-temporal regulation of cell migration^2^. Heparin, the most highly charged glycosaminoglycan in human ECM, is an important regulator of growth factor localization and retention due to its high binding affinity with many growth factors via electrostatic interactions, including growth factors that are important for wound repair (e.g. Vascular Endothelial Growth Factor [VEGF])^7–9^. Due to its high affinity for many growth factors, heparin has been incorporated homogeneously within biomaterial scaffolds to provide chemotactic bioactivity via controlled exogenous growth factor release^7,8^.

In addition to heparin-based controlled release of growth factors, multiple strategies have been developed to harness chemotactic gradients within biomaterial scaffolds, including those incorporating pre-programmed degradable carriers (e.g. GF-loaded PLGA nanoparticles) or advanced fabrication techniques (e.g. photopatterning)^10^. However, these techniques are limited by the macroscale (i.e. non-physiologic^2^) nature of their gradients^11^ and an inability to dynamically alter their signaling in response to tissue formation/remodeling. By contrast, in this manuscript we present an exciting heterogeneous heparin incorporation technique using spatially isolated hydrogel microspheres with covalently immobilized heparin, or heparin “micro-islands” (μIslands), that offer growth factor-mediated bioactivity, including chemotaxis, that is not reliant on exogenous growth factors. Our heparin μIslands create chemotactic gradients through reorganization of endogenous local growth factors.

To incorporate heparin μIslands in a format that allows cells to freely respond to chemotactic gradients, we took advantage of an injectable biomaterial platform, microporous annealed particle (MAP) scaffolds^12^. MAP scaffolds are composed of micron-scale spherical building blocks (microspheres) that can be mixed ratiometrically with heparin μIslands to achieve controlled heterogeneity. Further, MAP is assembled via covalent inter-microsphere bonding (i.e. annealing) *in situ* to form structurally stable scaffolds with cell-scale microporosity. In addition to observing the spontaneous reorganization of uniformly applied growth factors into micro-scale (i.e. physiologically relevant) gradients around heparin μIslands, we also validated their impact on cell chemotaxis and whole tissue regenerative behavior.

Specifically, we chose to focus the application of this phenomenon on diabetic wounds, which pathologically suffer from both a lack of organized tissue regeneration and growth factor retention^5,13,14^. Diabetic wounds result in approximately 130,000 lower limb amputations each year and affect 15% of diabetic patients in the U.S. (~10% of U.S. population has diabetes)^15,16^. Clinically, these wounds present a challenging healing environment that suffers, in part, from poor growth factor regulation^5,6,13,14^. Despite FDA approval of several therapies including decellularized matrices and recombinant growth factor (e.g. PDGF) delivery, their clinical use remains limited and even after treatment half of these wounds never heal^6,13,14,17^. We hypothesize that heparin μIslands, with their unique ability to concentrate growth factors into spontaneous microgradients, can improve the healing of these wounds. Therefore, we chose to test heparin μIslands in a relevant diabetic wound healing animal model^18^ using the most common class of advanced wound treatment (decellularized tissue^6,19^) as our clinically-relevant control.

### Particle building blocks to create a heterogeneous porous scaffold

Using our MAP scaffolds^12^ as a platform technology, we took advantage of the ability to mix and match particle populations to create ratiometrically-controlled heterogenous scaffolds while maintaining an injectable format. To design instructional interaction with chemokines, we chose to focus on heterogeneously distributed heparin particle populations (heparin μIslands). We hypothesized that heparin μIslands would locally sequester growth factors released endogenously in a wound environment and generate functional microscale gradients. Three particle populations with variable heparin concentrations (Hep_High_, Hep_Low_, and no Hep) were produced using a previously published high-throughput microfluidic method^20^ to isolate heparin concentration as the only changing variable between particle types by providing uniform geometric (diameter: 90μm, Supplemental Fig. 6) and mechanical (Young’s modulus: 18kPa, Supplemental Fig. 5) properties. The particles were composed of a synthetic hydrogel network of 4-arm poly(ethylene glycol) maleimide backbone crosslinked with a peptide sequence optimized for enzymatic resorption by matrix-metalloprotease-2 (MMP-2) and covalently bonded to an RGD cell adhesive peptide ligand (Fig. 1A). A custom heterofunctional 4-arm PEG maleimide/methacrylamide macromer was incorporated to facilitate scaffold annealing^21^. With consideration for our future diabetic wound healing assays, thiolated heparin^22^ was incorporated into the heparin μIslands at a concentration chosen to mimic mouse skin (Hep_High_) and one-tenth mouse skin (Hep_Low_) (Fig. 1B).

**Figure 1.**
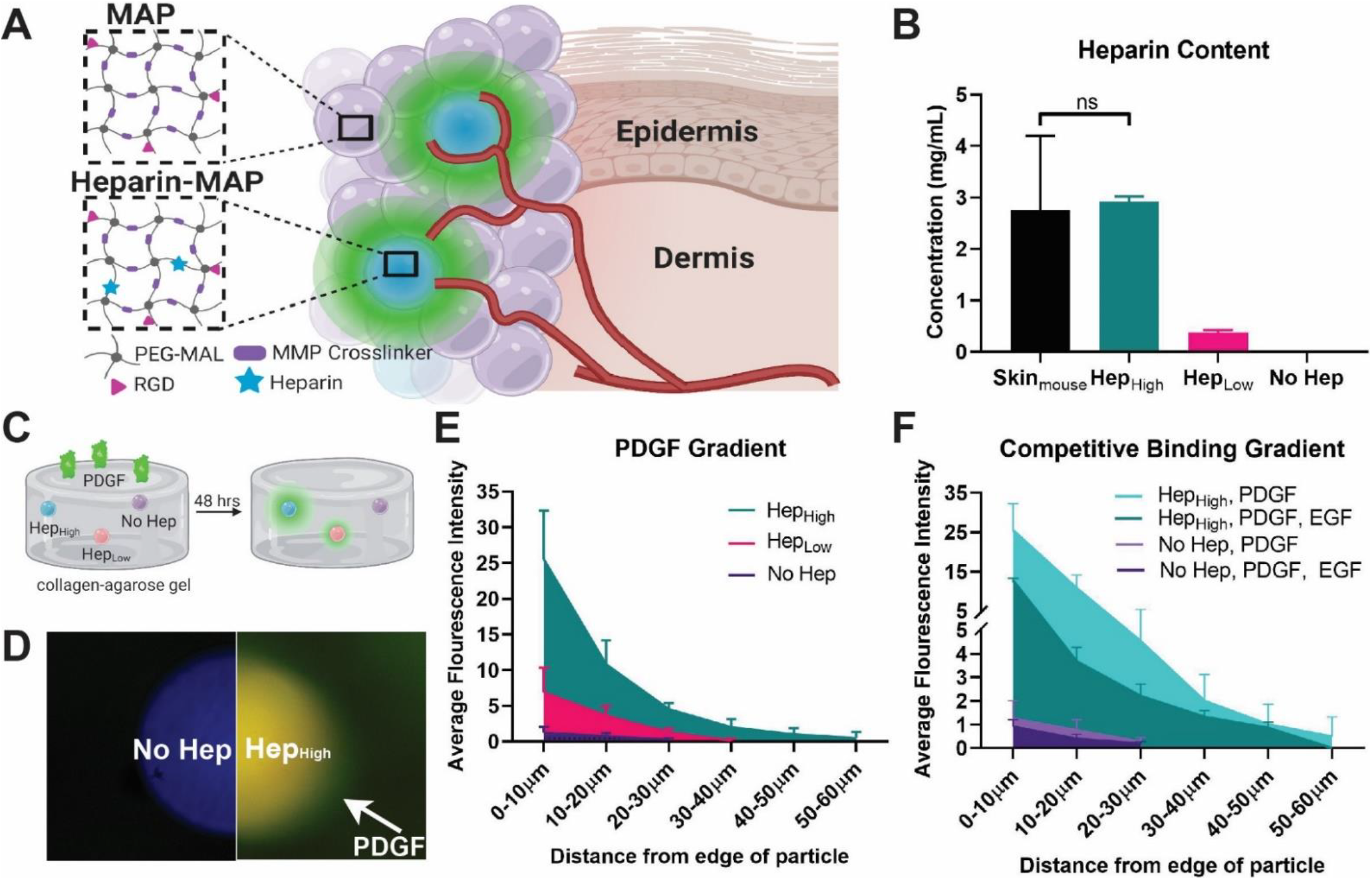
Synthesis and Gradient Characterization of Heparin μIslands. **A)** Particles were composed of a PEG-maleimide backbone and MMP-2 cleavable crosslinker with an RGD cell adhesive peptide with or without thiolated heparin. Small amounts of heparin particles (μIslands) were mixed with non-heparin particles to generate growth factor gradients. **B)** Heparin concentration within the particles was matched to mouse skin (Hep_High_) and one tenth of mouse skin (Hep_Low_). **C)** To test for gradient formation around our heparin μislands, we used a well-based assay with individual particles embedded within a collagen-agarose scaffold before introduction of a solution of biotin-labeled PDGF. **D)** After fixation, fluorescent Streptavidin revealed a PDGF gradient (green) around the heparin μIsland(red) and was absent around particle without heparin (blue). **E)** Quantification of fluorescence of concentric rings surrounding the microgels confirmed the presence of a gradient of PDGF. **F)** Gradient formation was maintained at a lower magnitude when microgels were incubated with PDGF and a competitive binding protein (EGF).

### Spontaneous microgradients from uniform growth factor

To validate our hypothesis that heparin μIslands can sequester and organize uniform growth factor distributions into microscale gradients, we embedded them within a collagen-agarose gel and incubated them with a solution of platelet-derived growth factor (PDGF) (Fig.1C). We chose PDGF for both its importance in wound healing and its known interactions with heparin^3,9^. After 48 hours of incubation with biotinylated PDGF, the gels were fixed and stained with fluorescent streptavidin. A gradient was visibly present around the heparin μIslands and was absent around particles without heparin (Fig. 1D). Quantification of relative gradient strength showed that increased levels of PDGF were detected out to 60μm around Hep_High_ μIslands and 40μm around Hep_Low_ μIslands (Fig. 1E). To test the impact of competitive binding on spontaneous gradient formation (a complication expected *in vivo*), the same PDGF assay was conducted again for the Hep_High_ particles with the additional complexity of added non-biotinylated Epidermal Growth Factor (EGF), which is also known for heparin-specific interactions. Spontaneous gradient formation was still observed around the Hep_High_ μIslands out to 60μm, but at lower magnitude than PDGF alone (Fig. 1F). Thus, with these *in vitro* assays, we were able to confirm an affinity-based structure-function relationship for heparin μIslands generating microscale gradients. These results prompted further investigation of potential effects on *in vitro* cell migration.

### Cell migration is dependent on particle spacing and heparin concentration

To further investigate the potential functional impact of heparin μIslands, we employed an *in vitro* spheroid migration assay developed for hydrogel constructs^23^ (Fig. 2B-C). Due to the potential for subsequent use in a diabetic dermal wound healing model, we chose two primary cell types relevant in dermal wound healing: human dermal fibroblasts and human dermal microvascular endothelial cells. Both particle ratio (heparin μIslands to no hep particles) and heparin concentration were varied to determine the effects of the heterogenous distribution of heparin μIslands (Fig. 2A). Notably, both ratio and concentration affected migration behavior for both cell types. Specifically, for ratios of heparin particles, we observed an interesting impact of heterogeneity, where just 1% Hep_High_ μIslands had no noticeable difference in migration compared to no heparin gels, but 10% had the greatest migration compared to all other groups (Fig. 2D-E). However, using just 10% Hep_Low_ particles only resulted in a slight increase in migration compared to no heparin (Fig. 2F-G), confirming that heparin concentration remains a critical factor. The cell migration in the 10% Hep_High_ μIslands was significantly greater than 100% Hep_Low_ which had the same total heparin in a homogenous format (Supplemental Figs. 9,10), verifying that heterogenous microenvironments promote the spatial cues necessary to increase migration without needing external factors (e.g. growth factors). The enhanced migration seen with 10% Hep_High_ μIslands in HDFs and HDMVECs prompted the use of this group for the diabetic wound healing studies.

**Figure 2.**
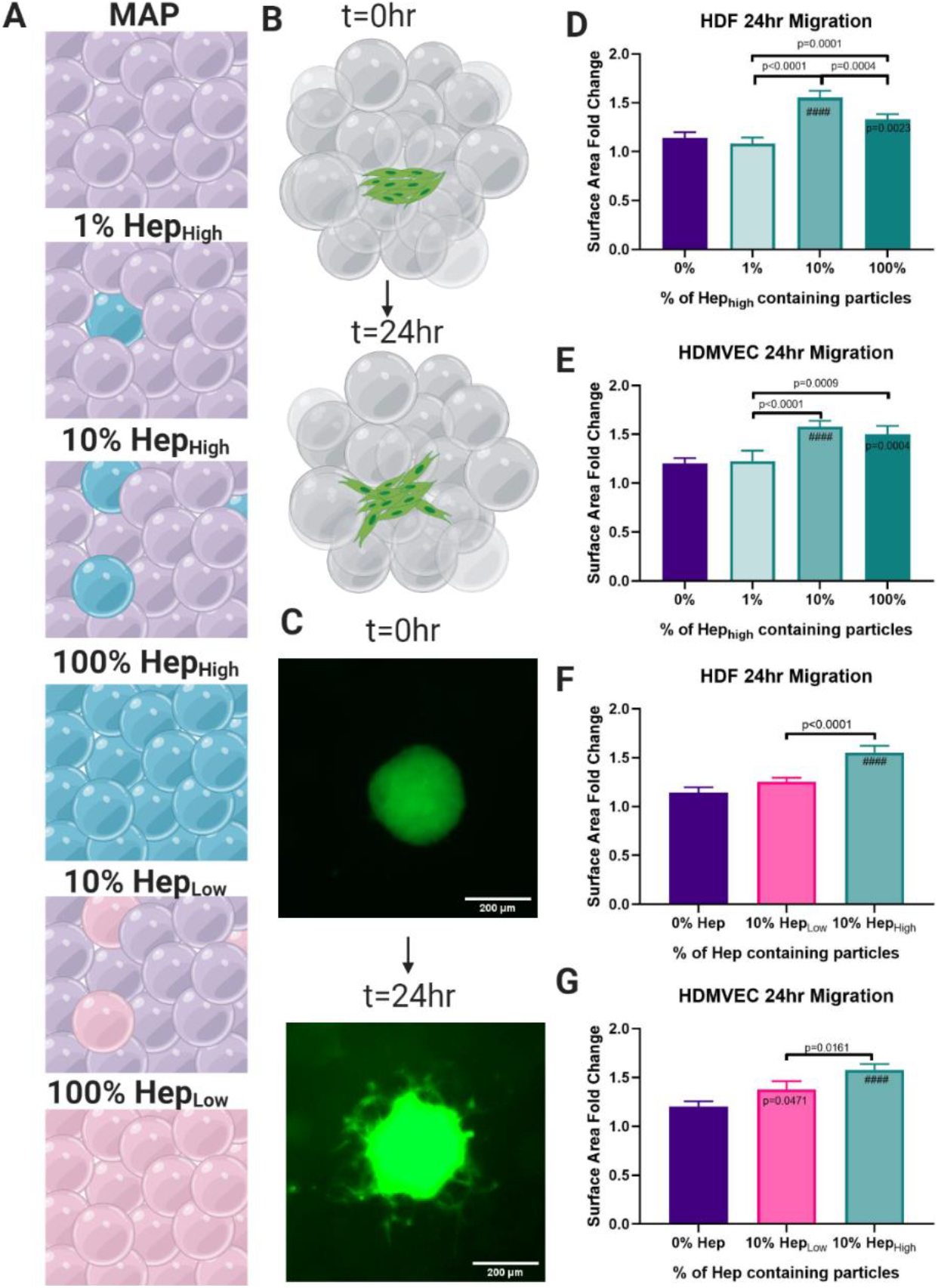
Cellular Migration. **A)** Six gel conditions were compared with varying concentrations and percentages of heparin particles. **B)** A spheroid migration assay was used to quantify cell migration over 24 hours. **C)** Representative images of spheroids at 0 and 24 hrs. **D)** Quantification of cell migration showed the 10% Hep_High_ μIslands exhibited the highest migration of Human Dermal Fibroblasts (HDFs) and **E)** Human Dermal Microvascular Endothelial Cells (HDMVECs) compared to all other groups with the same concentration of heparin, but different spacing. **F**) The 10% Hep_High_ μIslands significantly outperformed the 10% Hep_Low_ μIslands for HDFs and G) HDMVECs confirming the concentration of heparin and the spacing of heparin particles effect cell migration. Statistics: ANOVA, Multiple comparisons post-hoc tests (Tukey HSD). N=4. Significance inside of bars represents comparison to no hep gels. #### is p<0.0001.

### Heparin μIslands promote enhanced re-epithelialization in diabetic wounds

Diabetic wounds are characterized by their inability to move from the inflammatory to proliferation phase of wound healing, marked by the onset of re-epithelialization and re-vascularization. We chose two time points for this diabetic wound healing study, Day 3 and Day 7, to further characterize the progression of wound healing. The four treatment groups for this study were MAP gel with μIsands (10% Hep_High_), MAP gel without μIslands as a material platform control, Oasis wound matrix as an advanced clinical control (currently approved for use in diabetic wounds), and Aquaphor as a basic clinical control (OTC wound hydration product) (Fig. 3A). Quantification of wound re-epithelialization (determined by regenerated epidermal tissue or “tongues”) showed the μIslands group improved wound closure relative to all other groups (Fig. 3B,D). This data aligned with gross observations of wound granulation taken at Days 3 and 7 (Supplemental Fig. 12). By Day 7 the majority of wounds for all groups except Aquaphor had fully re-epithelialized the wound bed (Supplemental Fig. 13), as determined by Keratin-14 staining, and therefore epidermal thickness was quantified as a measure of dermal regeneration. The μIslands demonstrated a significantly greater epidermal thickness compared to the MAP and Oasis groups (Fig. 3C,D). In concert, these results demonstrate that the μIslands treatment provided a more regenerative and accelerated early healing result compared to all controls.

**Figure 3.**
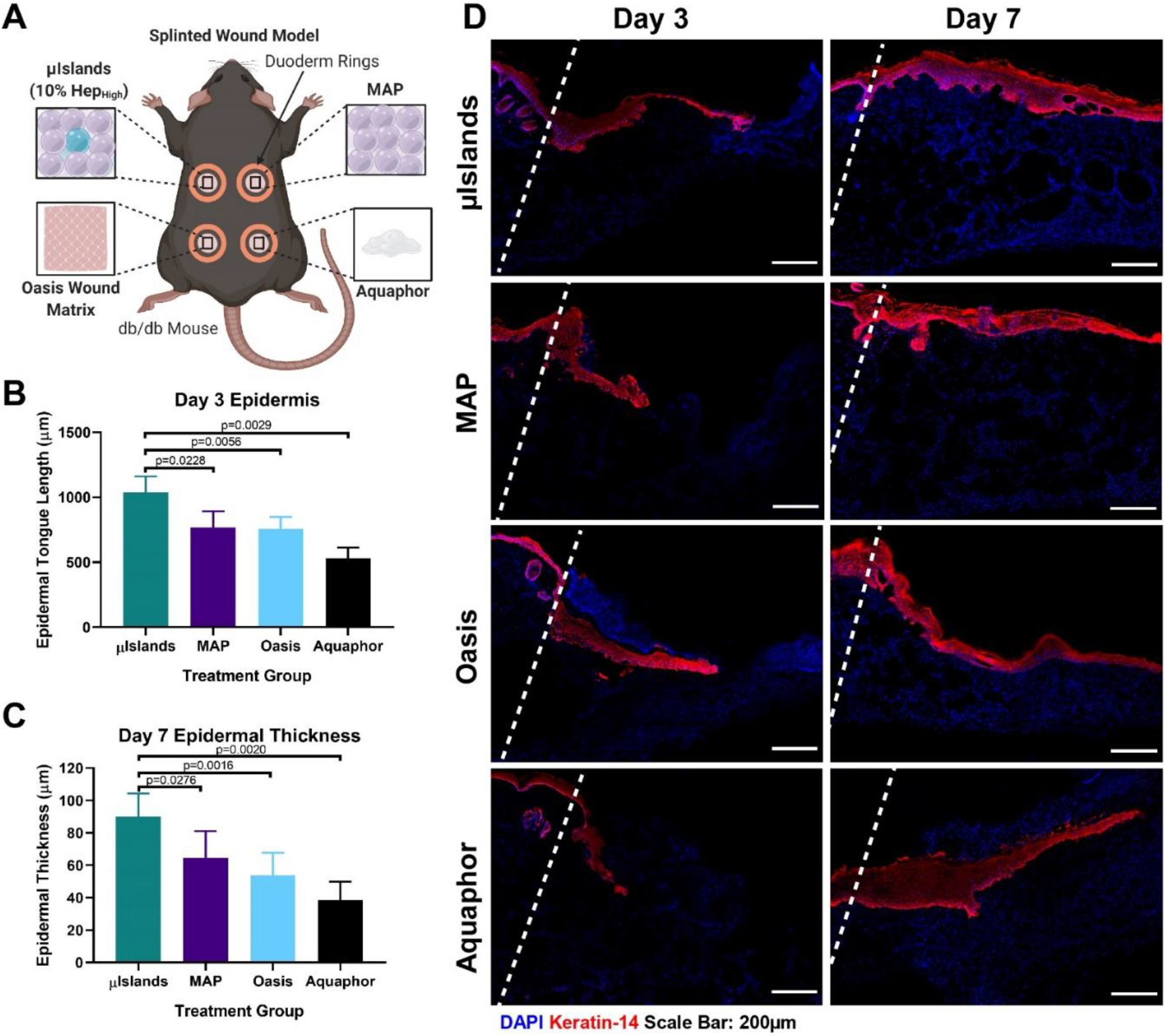
Epidermal Regeneration in a Diabetic Wound Healing Model. **A)** Four treatment conditions were evaluated in a mouse diabetic wound healing model at Day 3 and Day 7. **B)** Epidermal tongue length was quantified at Day 3 to determine extent of re-epithelialization. **C)** Epidermal thickness was quantified for all at Day 7 to compare stages of healing. **D)** Representative images of the four treatment groups keratin-14 staining at Day 3 and Day 7. Wound edges marked by dashed line. All graphs show mean +/− standard deviation. Statistics: ANOVA, Multiple comparisons post-hoc tests (Tukey HSD). N=6.

### Uniform revascularization at Day 7

Heparin’s known interaction with key growth factors in angiogenesis and the importance of re-vascularization to wound healing prompted us to investigate vascularization among the groups at Day 7. At Day 7, the μIslands group was characterized by the presence of extensive vasculature throughout the entire wound, while other groups had noticeably more of their re-vascularization occurring proximal to the wound edges (Fig. 4D). Quantification of this difference determined that μIslands provided significantly more staining overall for CD31^+^ endothelial cells and that the distribution of blood vessels was equal between the inner 50% of the wound and the edges (Fig. 4B). Interestingly, when analysis is limited to the edges of the wound, there were no significant differences in re-vascularization among MAP, Oasis, and the μIslands (Fig. 4B). By contrast, when analyzing the inner 50% of the wound, only the μIslands show as much re-vascularization as was present in the outer wound area. The Aquaphor group had minimal vessels observed within the wound. As an additional metric to characterize the new vasculature, we stained for pericytes (an indicator for vessel maturity) by staining for NG2^+^ cells proximal (<5 μm distant) to endothelial cells. Quantification of pericytes per wound area showed significantly more pericyte staining in the μIslands group, indicating more mature vasculature (Fig. 4C,E). In summary, μIslands produce a more mature and extensive vascular network throughout the entire diabetic wound by Day 7.

**Figure 4.**
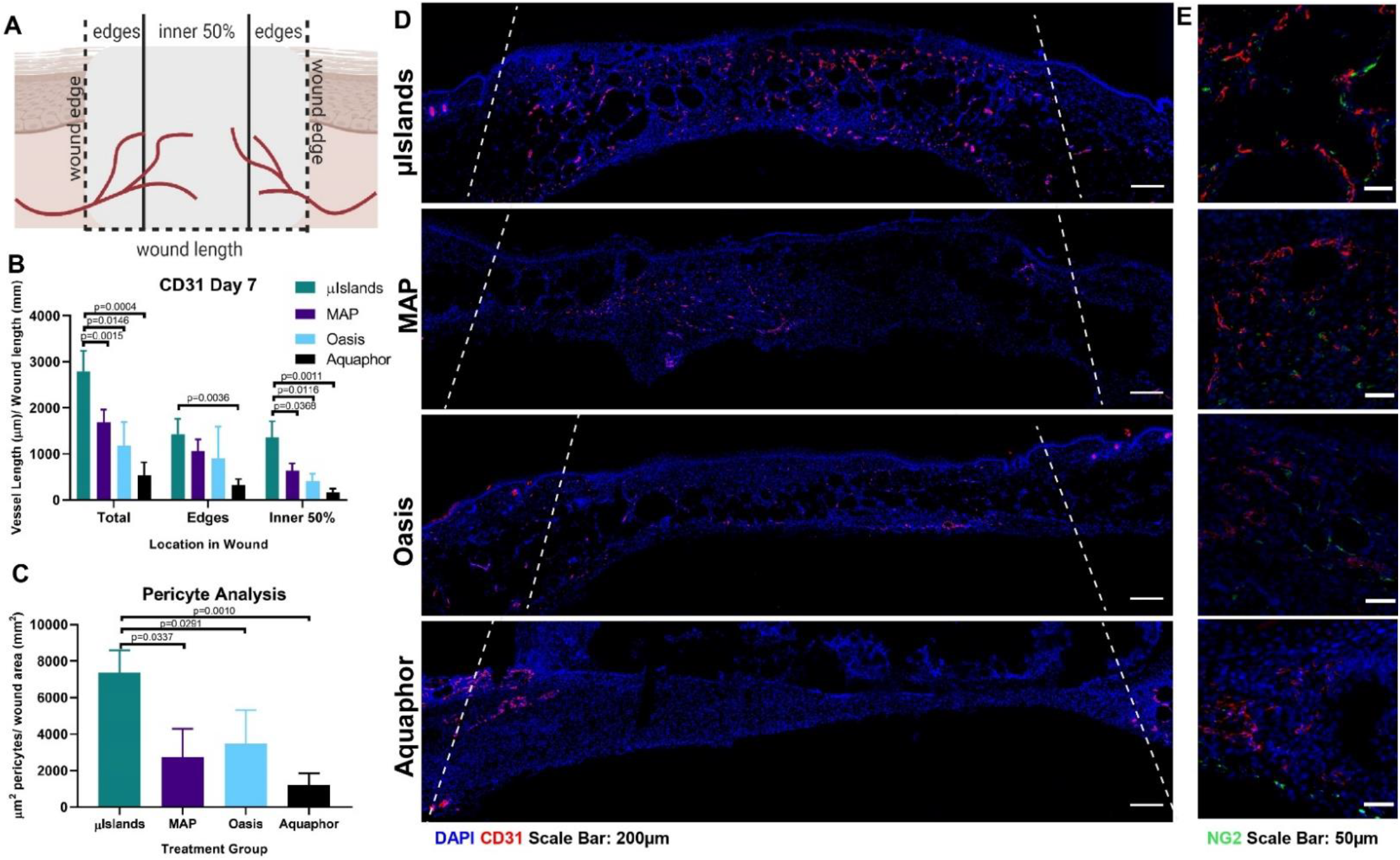
Vascularization at Day 7 in a Diabetic Wound Healing Model. **A)** Vessel analysis was performed to determine the extent of vascularization throughout the wound. **B)** Vessel length was quantified for the middle and edges of the wound at Day 7 as determined by CD31 staining. **C)** Pericyte analysis was performed to assess the presence of mature vasculature as indicated by proximal NG2 staining to a blood vessel. **D)** Representative images of the four treatment groups CD31 staining Day 7. Wound edges marked by dashed line. **E)** All graphs show mean +/− standard deviation. Statistics: ANOVA, Multiple comparisons post-hoc tests (Tukey HSD). N=6.

### No difference in immune modulation between μIslands and MAP

To investigate the potential for altered immune response caused by the presence of heparin μIslands, we stained for macrophage polarization at Days 3 and 7. There were no significant differences between any of the four groups for density of macrophages within the wounds (Supplemental Fig. 19). While we observed no difference between MAP with or without heparin μIslands, we did observe that MAP groups (with and without heparin μIslands) provided a clear immunomodulatory effect on wound macrophage polarization compared to either the Oasis or Aquaphor treatments (Fig. 5). Specifically, both MAP groups promoted more M2 than M1 polarization for both time points compared to Oasis and Aquaphor, and by Day 7 all macrophages were overwhelmingly M2 phenotype for both MAP conditions (Fig. 5A,C,D). Notably, these results aligned well with prior investigations of precast porous hydrogel scaffolds that are the geometric inverse of MAP gel^24,25^. Combined, these data demonstrate that the improved wound closure and re-vascularization effects observed for the heparin μIslands group was likely not due to changes to the immune response within the wound.

**Figure 5.**
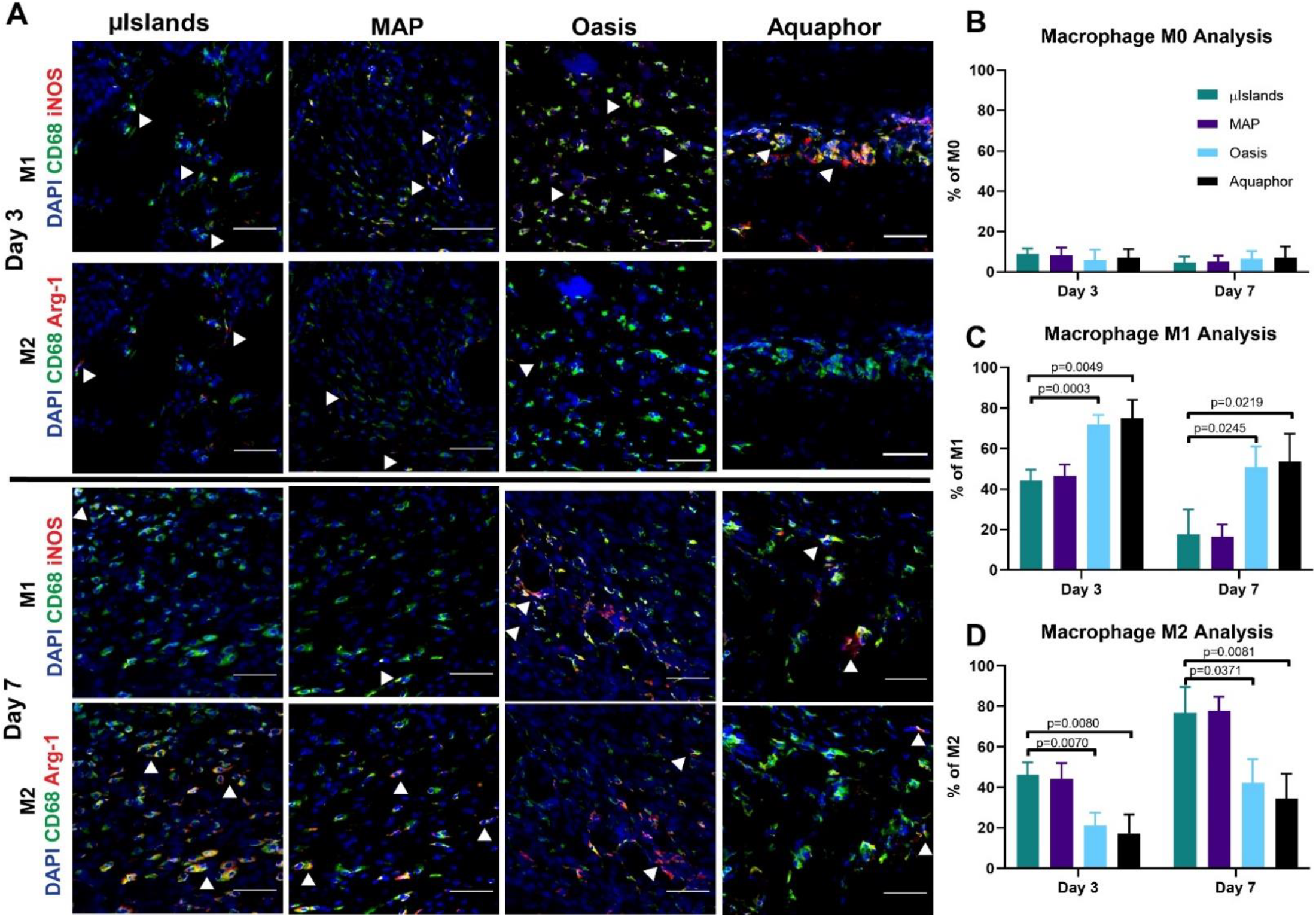
Macrophage Polarization. **A)** Representative images at Day 3 and Day 7 of macrophage M1 and M2 polarizations. **B)** Percentage of macrophages in the wounds for each group at Day 3 and 7 that only stained positive for CD68. **C)** Percentage of macrophages in the wounds that stained positive for the M1 inflammatory phenotype as indicated by CD68+ and iNOS+. **D)** Percentage of macrophages in the wounds that stained positive for the M2 pro-regenerative phenotype as indicated by CD68+ and Arg-1+. Statistics: ANOVA, Multiple comparisons post-hoc tests (Tukey HSD). N=6.

## Conclusions

Here we present a new type of bioactive scaffold which can harness local growth factors to form microscale chemotactic gradients, eliminating the need for exogenous delivery while maintaining an instructional microenvironment. Previously, creating gradients at this scale within biomaterials has required advanced biofabrication strategies (photolithography, bioprinting) that limits injectability as well as the ability to fill large wounds. MAP with heparin μIslands significantly improved diabetic wound healing outcomes compared to an unaltered MAP gel and two clinically relevant control groups. We plan to use future studies to further understand the *in vivo* mechanism of improved healing by identifying and quantifying the cytokines being organized by the bioactive heparin μIslands. The presented approach has high translational potential for applications requiring quick tissue integration in challenging tissue environments.

## Acknowledgements

Figure schematics created with BioRender.com. LP was supported by a National Science Foundation Graduate Research Fellowship and by the National Heart, Lung, and Blood Institute of the National Institutes of Health under Award Number F31HL154731. This work was partially supported through the US National Institutes of Health High Priority, Short-Term Project Award (1R56DK126020-01) and The Wallace H. Coulter Translational Partners Program at The University of Virginia.

## Notes

### Competing Interest Statement

D.R.G. has financial interests in Tempo Therapeutics which aims to commercialize MAP technology.

### Summary of Updates

Slight alteration to manuscript title and revision of a figure scheme mistake (removal of misplaced single pink sphere from Figure 2a -- 100% HepHigh).

