## Supplemental Information for "Heparin MicroIslands to Promote Enhanced Diabetic Wound Healing Outcomes"

**Supplementary Information**

### Methods

*Biomaterial Creation and Characterization:*
*Sources/ Storage of Materials:* Four-arm poly(ethylene glycol) maleimide (PEG-MAL, 10kDa and 20kDa) was purchased from Nippon Oil Foundry, Inc (Japan). RGD cell adhesive peptide (Ac-RGDSPGGC-$NH_2$) and the MMP-2 degradable crosslinker (Ac-GCGPQGIAGQDGCG-$NH_2$) were purchased from WatsonBio Sciences. All materials were dissolved in either ultrapure water or 0.1% Trifluoracetic Acid solutions (to prevent disulfide bond formation) and aliquoted into specified amounts to ensure precision. MethMal was synthesized as previously reported[21]. The aliquots were lyophilized and stored in -20°C until preparing aqueous gel solutions.
*Thiolation of Heparin:* Heparin (MW=15,000Da, Millipore Sigma) was dissolved at 20mg/mL in ultrapure water (300mg reactions, N=3). A 100mM (3-(2-pyridyldithiol) propionyl hydrazide) (PDPH) (CovaChem) solution was prepared in ultrapure water. The 100mM stock PDPH solution was added to each heparin reaction to target a 20% modification. 6.894 ul of AlexaFlour hydrazide 555 (1mg/mL) was added to each reaction to fluorescently tag the molecule. 145µLof the 100mM PDPH stock solution was added to each 300mg reaction of heparin. The reactions were mixed well and then the pH was adjusted to 6.5. DMTMM was added to each reaction in a 1:1 molar ratio to the heparin repeat units (estimate 619.49 Da[26]) each day of the reaction. The reaction was allowed to proceed for 3 days on a rotator at room temperature. Each reaction was dialyzed using 3.5kDa snakeskin dialysis tubing for 3 days in 1M NaCl (4L, changed 2x daily) and then 0.01M NaCl (4L, 6x1hr washes). The product was frozen and lyophilized. The volume before and after dialysis were recorded to determine the percentage of mass after lyophilization that is salt to give a purity value. After quantification of heparin thiolation, the three batches were combined and deprotected with 25mM TCEP (Sigma) at room temperature for 15 minutes. The deprotected solution was dialyzed using 3.5kDa snakeskin dialysis tubing (Fisher) in 0.01% trifluoracetic acid and 1M NaCl (4L, 1x3hrs), then 0.1M NaCl (4L, 2x1hrs), then 0.01M NaCl (4L, 2x1hr). The product was frozen and lyophilized, then stored at -20°C until use.
*Quantification of heparin thiolation:* Quantification of thiol concentration was based on a deprotection assay following PDPH manufacturers protocol and the modification was verified by NMR. The Thermofisher Scientific protocol for Pyridine-2-Thione Assay was used to determine the level of sulfhydryl modification. Briefly, the absorbance of the modified heparin was measured using a nanodrop before and after exposure to 1mg/mL DTT for 15 min at 343nm (Supplemental Fig. 1B). HNMR was also used to confirm PDPH modification. A Varian Inova 500 NMR spectrometer located in the UVA Biomolecular Magnetic Resonance Facility was used to acquire the spectra. 25 mg of each reaction was dissolved in deuterium oxide (Sigma). MestReNova was used for analysis, and 3 PDPH peaks (~7.4, 7.95, 8.45ppm) were compared to the acetyl peak of heparin(~2.1ppm) (Supplemental Fig. 1C).
*Gel Formulations:*
*No Hep formulation:* A 3.2wt% (w/v) gel was used for this formulation. The final concentrations in the gel were 45.88mg/mL PEG-MAL (20kDa), 0.82 mg/mL RGD, 8.06mg/mL MethMal, and 4.62mg/mL MMP-2 crosslinker. For macrogel formulations, the PEG-MAL, RGD and MethMal were dissolved in a pH=4.5 10X PBS solution and the MMP-2 crosslinker and heparin were dissolved in pH=7.4 10X PBS. For microgel synthesis the PEG-MAL, RGD, and MethMal were dissolved in pH=1.5 10X PBS and the MMP-2 crosslinker along with 5µM of biotin labeled maleimide was dissolved in pH=7.4 1X PBS.
*$Hep_{High}$ formulation:* A 2.2wt% (w/v) 6mg/mL heparin gel was used for this formulation. The final concentrations in the gel were 34.83mg/mL PEG-MAL (10kDa), 0.82 mg/mL RGD, 8.06mg/mL MethMal, 5.56mg/mL MMP-2 crosslinker, and 6mg/mL heparin. For macrogel formulations, the PEG-MAL, RGD and MethMal were dissolved in a pH=4.5 10X PBS solution and the MMP-2 crosslinker and heparin were dissolved in pH=7.4 10X PBS. For microgel synthesis the PEG-MAL, RGD, and MethMal were dissolved in pH=1.5 10X PBS and the heparin, 5µM of Alexa Fluor labeled maleimide, and MMP-2 crosslinker was dissolved in pH=7.4 1X PBS.

*$Hep_{Low}$ formulation:* A 3.0wt% (w/v) 0.6mg/mL heparin gel was used for this formulation. The final concentrations in the gel were 38.89mg/mL PEG-MAL (10kDa), 0.82 mg/mL RGD, 8.06mg/mL MethMal, 7.47mg/mL MMP-2 crosslinker, and 0.6mg/mL heparin. For macrogel formulations, the PEG-MAL, RGD and MethMal were dissolved in a pH=4.5 10X PBS solution and the MMP-2 crosslinker and heparin were dissolved in pH=7.4 10X PBS. For microgel synthesis the PEG-MAL, RGD, and MethMal were dissolved in pH=1.5 10X PBS and the heparin, 5µM of Alexa-Fluor labeled maleimide, and MMP-2 crosslinker was dissolved in pH=7.4 1X PBS.

*Gelation kinetics:* Macro-scale gels (i.e. macrogels) of 400µL volume were formed on a ThermoScientific Viscometer (20 mm plate attachment). Sufficient gelation time was determined by a minimal change (<10%) in storage modulus over 15 minutes using HAAKE RheoWin (Supplemental Fig. 3).

*Macrogel production:* Macro-scale gels (i.e. macrogels) were used to mechanically match hydrogel stiffnesses between groups to be approximately 15-20kPa. 200µL gel pucks were formed between SigmaCote® coated slides for 30 min past total gelation according to the viscometer. The macrogels were collected and swollen to equilibrium in PBS overnight at 37°C prior to testing.

*Microgel production:* Microgels were produced using a PDMS mold for 45-55µm particles described in Rutte, et. al[20]. Picosurf surfactant (Sphere Fluidics) was diluted to a 1% solution in NOVEC 7500 (3M). The gel solutions were prepared as described above, however a 10X PBS solution pH=1.5 was used to dissolve the backbone solution to ensure gelation would not occur in the device. Using a syringe pump, the surfactant and aqueous solutions were run through the device at 5mL/ hour and collected in a 50mL conical tube. The microgels were mixed with a solution of triethylamine (20µl/mL of gel) diluted in NOVEC 7500 to ensure complete gelation before purification.

*Microgel purification:* Microgels were washed three times with NOVEC 7500 (1X gel volume). Next, microgels were swelled in PBS (5X volume) and washed 3 more times with NOVEC 7500, allowing separation of the aqueous and oil solutions via settling. The NOVEC oil was removed and microgels were washed with PBS (5X gel volume) and Hexanes (5X gel volume) and centrifuged at 4696gx5min. Particles were quenched with a 100mM N-Acetyl-L-Cysteine (Acros Organics) solution in PBS overnight at 37 deg C to quench excess maleimides.

*Microgel sterilization:* All remaining steps were performed in a biosafety cabinet. After quenching, microgels were washed three times with 70% isopropanol (5X gel volume, 4696gx5min). Microgels were stored at 4 deg C in 70% isopropanol until being used.

*Materials Characterization Assays:*

*Heparin Quantification in Skin:* Skin of Swiss Webster mice was explanted and digested to isolate heparin using an adaptation of the protocol from Zuo, et. al[27]. Briefly, the tissue specimens are extracted and digested with proteinase K. The amount of total sulfated GAG is quantified using a dimethylmethylene blue (DMMB) assay[28] (Supplemental Fig. 2). Following a protocol by Hahn, et. al[29], the heparin proportion of GAGs was determined by the loss of DMMB signal after sample exposure to heparinase following manufacturer's protocol (R&D Biosystems).

*Heparin Incorporation in Macrogels:* Heparin was tagged with AF555 during the PDPH modification. A standard curve of heparin fluorescence (Supplemental Fig. 4A-B) was produced by making serial dilutions of the heparin dissolved in PBS. 150µl of each solution was placed in a 96 well Greiner SensoPlate and imaged using an ImageXpress MicroConfocal Imaging System (Molecular Devices). A 5mm biopsy punch of each macrogel containing heparin was placed into a well of a 96 well Greiner SensoPlate and covered with 100ul of PBS. Using the same focus and exposure time as the standard curve the macrogel is acquired. Using a custom ImageXpress module (Molecular Devices) the average intensities for each site of the macrogel are calculated (Supplemental Fig. 4C). The 4 sites with the smallest standard deviation are averaged and the concentration is calculated based on the standard curve (Supplemental Fig. 4D).

*Mechanical Analysis of Macrogels:* Macrogels were removed from PBS and excess moisture was wicked from the pucks before testing. An Instron mechanical load device was used to test compressive stiffness (Young's modulus) at a rate of 0.5mm/min for 1 mm (Supplemental Fig. 5A) and BlueHill® software analyzed the load (N) and extension (mm). Stress-strain curves were produced and the Young's modulus (Pa) was calculated using the MATLAB SLM package (Supplemental Fig. 5B).

*Particle Size Characterization:* Microgel particle spherical diameter was determined using fluorescent images of a dilute solution (1:100) of microspheres and ImageXpress based quantification (Molecular Devices). A minimum of 500 particles (N) were analyzed to determine average diameter (D) and polydispersity index (PDI). The number average was calculated by $Dn = \sum NiDi / \sum Ni$ and weight average was calculated by $Dw = \sum NiDi^2 / \sum NiDi$. PDI was calculated by Dw/Dn. (Supplemental Fig. 6)

***In vitro studies:***

*Gradient Formation:* Platelet Derived Growth Factor (PDGF, R&D Biosystems) was biotinylated (NHS Sulfo EZ Link Biotin) according to manufacturer's protocol to allow for post-fixation identification (Fisher). MAP particles (no hep, $Hep_{Low}$, and $Hep_{High}$) were embedded (1:100) in 100ul collagen-agarose gel (1mg/mL collagen, 1% agarose) for structural support. Particles were labeled with different Alexa Flour maleimide tags to identify the different types. After gelation at 37 deg C, the gels were soaked in 10% FBS for 8 hours to mimic *in situ* conditions. Finally, the solutions were removed and then incubated with the growth factor of interested at 1ug/mL in 1% BSA for 48 hours on a shaker plate at 37 deg C. Following incubation, the gels were frozen in OCT and then sectioned into 10µm sections using a cryostat. To visualize the gradients, the slides were rehydrated with PBS and then stained with streptavidin labeled with Cy5 (1:200 dilution) for 1 hour at room temperature. Slides were washed three times with PBS and then coverslipped and imaged at 20x using an ImageXpress Micro-Confocal (Molecular Devices).

*Image Analysis:* ImageJ based analysis was used to calculate the gradient formation. The particles were traced using ImageJ in the fluorescence channel that corresponded to the specific particle type. That tracing was overlayed on the Streptavidin fluorescent channel corresponding to the PDGF. 10µm concentric rings were generated surrounding the particle tracing and the mean gray value was measured using ImageJ (Supplemental Fig. 7). All values were subtracted from the background mean gray value in an area without a staining to generate the value above background fluorescence.

*Sources of Cells:* Primary adult Human Dermal Fibroblasts (HDFas) and primary Human Dermal Microvascular Endothelial Cells (HDMVECs) were purchased from ATCC and cultured according to manufacturer's guidelines.

*Cell Migration- Spheroid Formation:* Before starting the study, HDFs and HDMVECs were fluorescently labeled using AF488 CellTracker following manufacturer's guidelines (Fisher). Spheroids were formed at a concentration of 50,000 cells/mL for HDMVECs and 125,000 cells/mL for HDFs. Methylcellulose was prepared following a previously published protocol[30], and it was supplemented as 20% of the media to create circular spheroids. 20ul spheroids were incubated using the hanging drop method for 48 hours, and spheroid formation was confirmed using a brightfield microscope (Supplemental Fig. 8).

*Cell Migration Assay:* Cell migration was performed using an assay previously developed to measure migration in hydrogel scaffolds[23]. MAP gel formulations were incubated in complete cell culture media at 37°C overnight before starting the study. After incubation, microgels were spun down at 4696gx5min and suspended in a 2mM LAP (Sigma) solution in media at a 1:1 concentration. Microgels were spun down again at a high speed and the excess solution was poured off. The groups for each study were 1% heparin high, 10% heparin high, 100% heparin high, 10% heparin low, 100% heparin low and no heparin gel. 50µl pucks of gel were annealed using UV light at 14.8mW/cm$^2$ between sterile Sigma-coated slides for 30 seconds and then transferred to the bottom of a 24 well plate. Spheroids were pipetted onto the top of the gel and then imaged to get the 0- hour time point. At 24 hours the spheroids were imaged again using the ImageXpress Micro-Confocal (Molecular Devices). Fold change was measured using ImageJ as previously described[23] using Fiji.

<u>*In Vivo Studies:*</u>

*Mouse Excisional Wound Healing Model*: All animal surgeries were performed in accordance with Animal Care and Use Committee guidelines under UVA Animal Protocol 4165. 8-week-old male db/db mice (Jackson stock: 000697 HOM 1) were used for all studies. Mouse excisional wound healing experiments were performed based on a previous protocol for mouse wound healing favoring cutaneous regeneration and preventing wound contraction by splinting[31].

*Excisional Wound Healing Model Surgery*: Mice were anesthetized using continuous application of isoflurane throughout the duration of the procedure. Hair was removed from the dorsal side of the mouse the day prior to surgery using a combination of electric clippers and Nair, followed by PBS washes to remove the hair. Nails were trimmed to lessen the chance of splint removal. The day of surgery, the mouse backs were first sterilized with iodine and 70% isopropanol. A sterile 5mm biopsy punch was used to create 4 symmetrical full thickness wounds on the back, two on each side of the midline. Surgical scissors were used to cut out the wound from the biopsy punch indention. Duoderm rings stacked 4 high were used to create the splints. Prior to surgery they were prepared by stacking four layers on top of one another and using a 12mm biopsy punch to create the outside of the splint and 8mm biopsy punch to create the inside of the splint. Duoderm splints were superglued (Gorilla Glue) around the wound to prevent contraction. Next, each treatment was applied to the wound in a randomized order which was rotated for each mouse. For MAP conditions, 12µL of gel was applied and smoothed into a flat surface using a positive displacement pipette. Scaffolds were annealed for 30 seconds using 365nm UV light (15.4cm away, 440mA, Thor Labs). Oasis was prepared by using a 5mm biopsy punch to cut out the matrix and then placing it directly on the wound. Aquaphor was spread on the wound with a sterile cotton swab. Pictures of each wound were taken after the treatment was applied. Finally, wounds were covered with small squares of Telfa pads which were superglued on the duoderm rings to protect the wounds. Mice received 0.1cc of buprinex immediately following surgery and 24 hours later. Mice were singly housed and checked twice a day to confirm they had not removed their bandages. At the end of the wound healing experiments (3 days for six mice, 7 days for six mice) the wounds were imaged again. Mice were sacrificed by isoflurane overdose and cervical dislocation. The wound samples were retrieved with surgical scissors and immediately embedded in OCT and frozen with dry ice for cryosectioning.

*Evaluation of Wound Closure:* Wounds were imaged on day 0 and either day 3 or 7 for each mouse. Closure fraction was determined by comparing the pixel area of the wound to the pixel area within the duoderm ring (Supplemental Fig. 11). Closure fractions were normalized to Day 0 for each condition. Wound closure analysis was performed with investigators blinded to the treatment group identity during analysis. Any wound that had the bandage removed at some point during the study by the mouse was removed from this analysis due to the potential for contraction.

*Tissue Section Immunofluorescence:* OCT blocks were sectioned with 20µm sections with 2 sections at different locations in the wound on each slide. Slides containing tissue sections were stored at -80 deg C until staining. Slides containing tissue sections were first fixed in acetone for 10 min. Slides were then washed with PBS (3 x 5min) and then blocked with 1X PBS, 5% goat serum, 0.1% Triton-X and 1% FC block. Following blocking, the slides were incubated with the primary antibody prepared in 5% goat serum listed in Supplementary Table 1 overnight at 4 deg C. Secondary antibodies were all prepared in 1X PBS at a 1:1000 dilution. Following incubations slides were washed in PBS (3x5min) to remove any primary antibody and then incubated with the secondary antibody for 1 hour at RT. Following secondary incubation slides were washed again with PBS. Next the slides were stained with DAPI at a 1:1000 dilution in PBS for 20min at RT. Slides were washed again and then mounted with Prolong Gold Antifade mounting medium and stored at 4 deg C before being imaged.

*Keratin-14 Analysis- Epidermal Tongue Length:* Day 3 wound sections were stained with keratin-14 and imaged at 10X using an EVOS microscope. Epidermal tongue length was traced on at least one side of the wound for two sections at different locations within the wound. Tongue length was measured on ImageJ using the line tool and tracing the top of the epidermal tongue from where it starts to ends and measuring the length (Supplemental Fig. 14). The wound edge was determined by the change in morphology from the surrounding tissue typically indicated by an increase in thickness, slope downward, or lack of hair follicles.

*Keratin-14 Analysis – Epidermal Thickness:* Epidermal thickness was quantified for wounds at Day 7. If a wound was not fully re-epithelialized, the thickness of the epidermal tongue was quantified. Epidermal thickness was quantified by tracing the area of keratin staining from the wound edges on ImageJ to calculate an area and then dividing it by the length of the wound, which was also measured using the ImageJ (Supplemental Fig. 15). Epidermal thickness was quantified for two sections from each wound.

*CD31 Analysis:* Full wounds were imaged at 10X objective using a Molecular Devices Confocal Microscope to visualize CD31 staining and DAPI. Images were cropped to only contain the wound region and not surrounding tissue for quantification. Images were uploaded into the MATLAB REAVER GUI[32] and set to analyze all wounds for vasculature with settings of 400 as the minimum connected component area and vessel thickness set to 1. This batch analysis exported the vessel length and vessel area (Supplemental Fig. 16). These measurements were normalized by tracing the wound length and area in ImageJ. Additionally, a binary image of each quantification was exported. From this image we were able to export X,Y coordinates of each vessel location and determine the percentage of vessels in each quarter of the wound. Vascularization was quantified for two full sections from each wound.

*Pericyte Analysis*: Full wounds were imaged at 20X objective using a Molecular Devices Confocal Microscope to visualize CD31, NG2, PDGFRB, and DAPI stains. Images were cropped to only contain the wound region and were analyzed for vessels as described above. The binary image of the vessels was overlayed with the NG2 channel from the original image using ImageJ. NG2 positive cells were cropped out of the image if they were not determined to be proximal to the endothelial cells (<5µm). The area of the NG2 positive staining that was determined to be proximal to vessels was calculated in ImageJ and normalized to the wound area (Supplemental Fig. 17).

*Macrophage Analysis:* Full wounds were imaged at 20X objective using a Molecular Devices MicroConfocal Microscope to visualize DAPI, CD68, Arg-1, and iNOS staining. Stitched sections were cropped to only the wound bed and not surrounding skin. After cropping, background removal was performed using ImageJ and each channel was thresholded with the Moments filter to obtain positive staining. Thresholded images were read into MATLAB to determine co-localization (Supplemental Fig. 18). Macrophages were determined to be M1 if there was positive staining for DAPI, CD68, and iNOS. Macrophages were determined to be M2 if there was co-localized positive staining for DAPI, CD68, and Arg-1. Macrophages that only stained positive for DAPI and CD68 were M0. Macrophage density was determined by calculating the number of total macrophages divided by the wound area. Macrophage polarization was quantified for two full sections from each wound.

#### Supplemental Figures

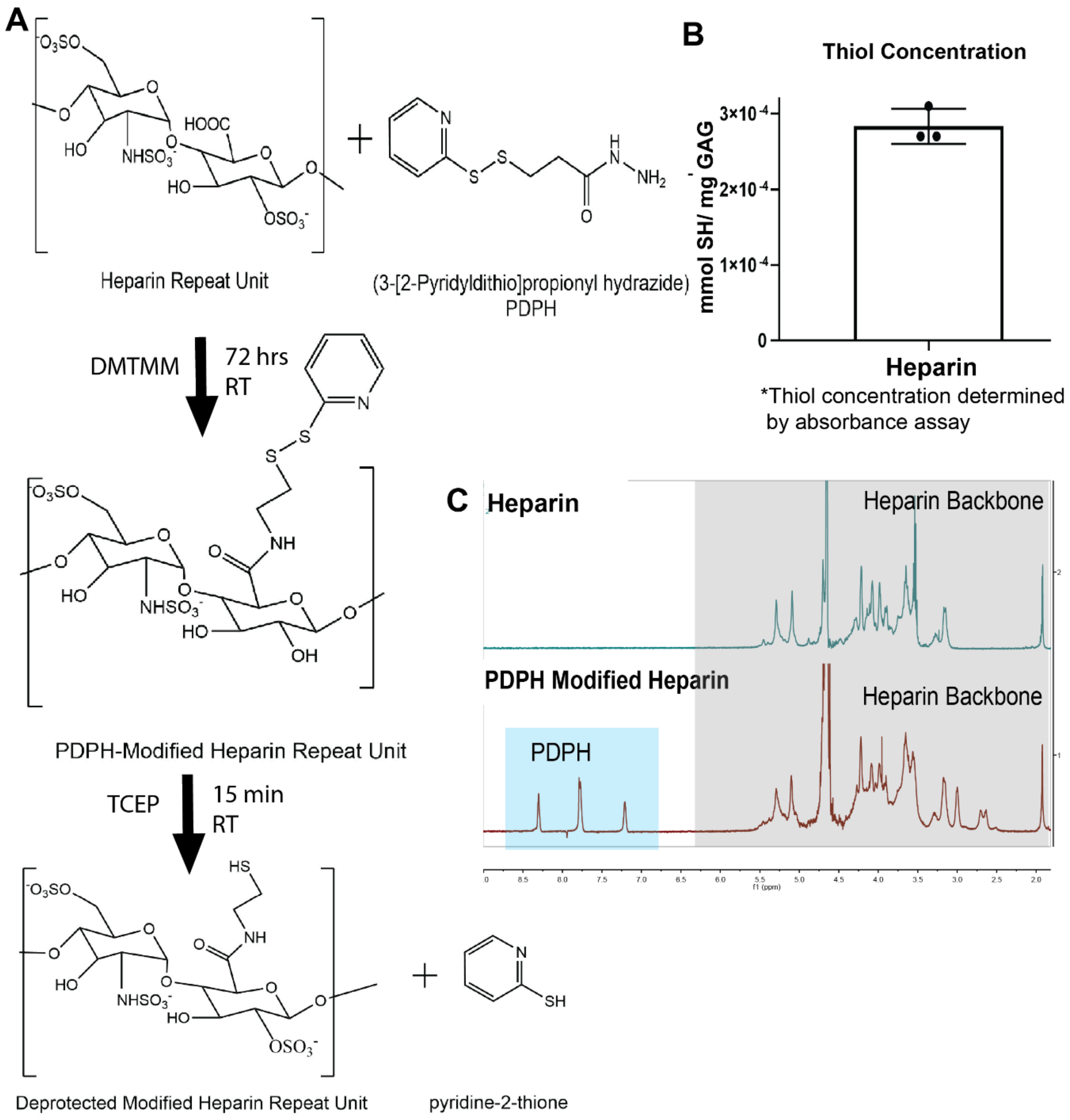


**Supplemental Figure 1: A)** Heparin was thiolated with PDPH (3-(2-pyridyldithio)propionyl hydrazide) and flourescently labeled (Alexa flour 555-hydrazide). Thiolation extent was determined via a **B)** deprotection absorbance assay and **C)** confirmed via HNMR.

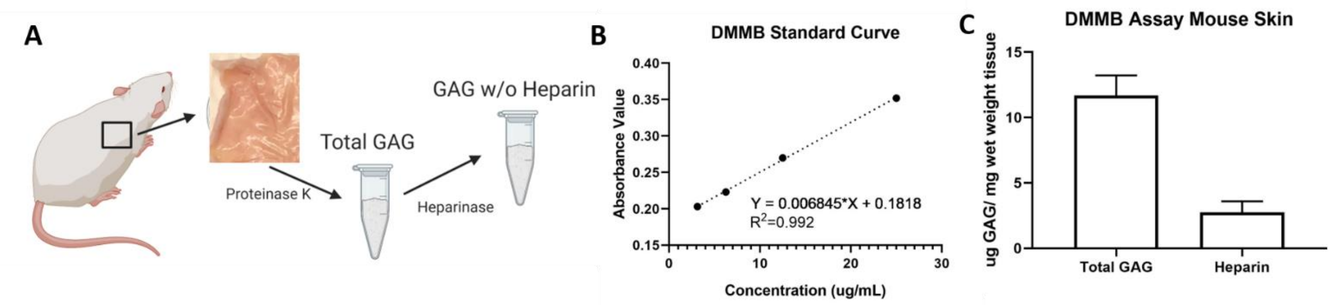


**Supplemental Figure 2: A)** Swiss webster mouse skin was digested with proteinase K to determine total GAG and with heparinase . **B)** A dimethylmethylene blue (DMMB) assay was performed on digested skin both before and after heparinase digestion to **C)** calculate quantify GAG concentration and heparin concentration. Graphs represent mean+/- standard deviation. Statistics: ANOVA, N=3.

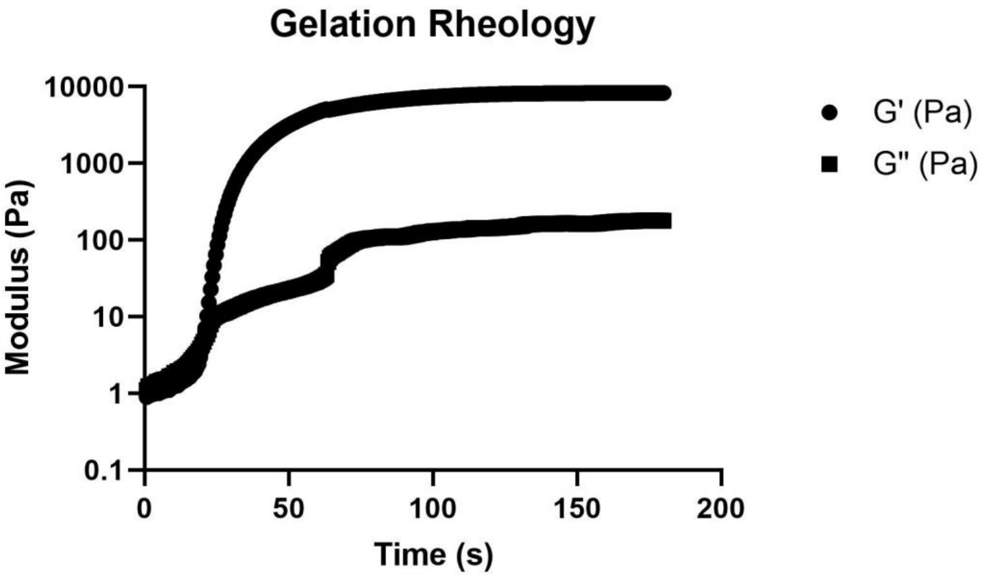


**Supplemental Figure 3:** A representative curve of the rheologic analysis used to ensure full gelation of macrogels prior to swelling.

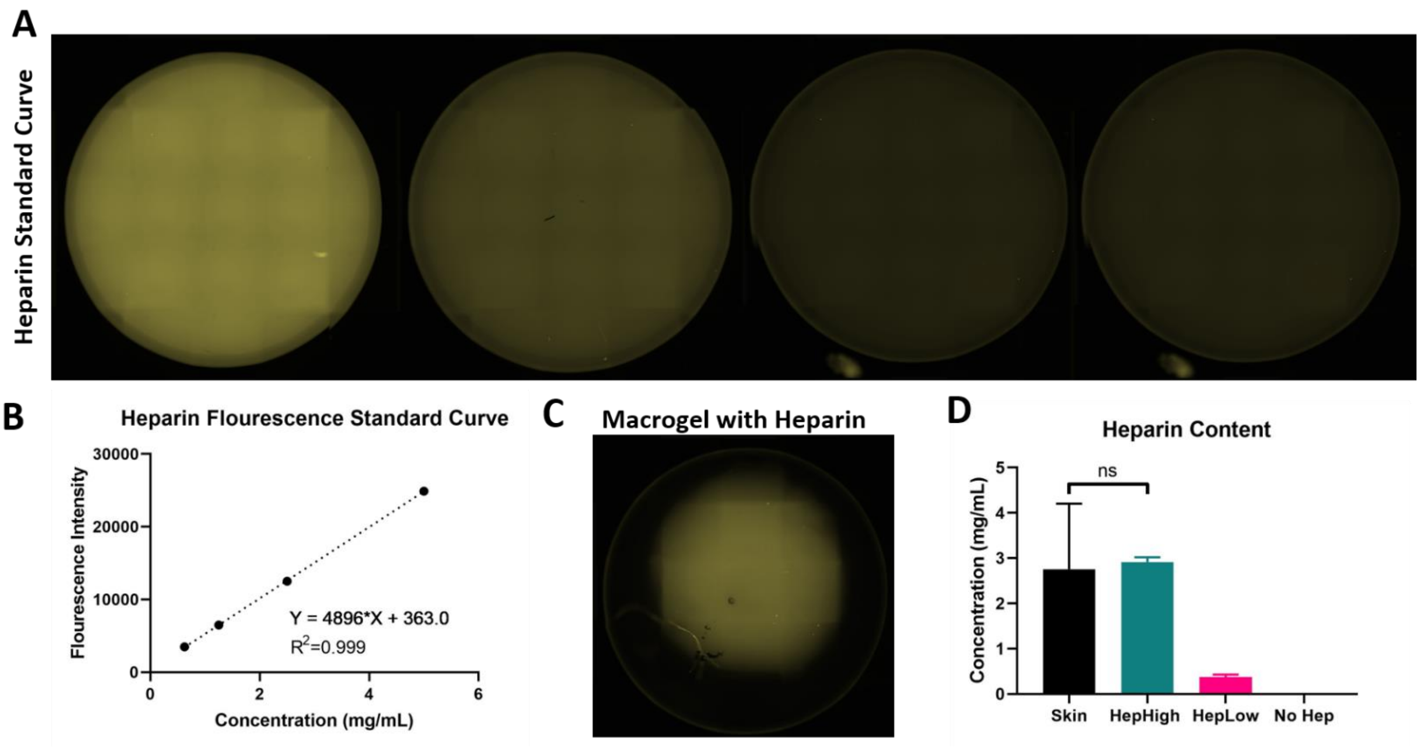


**Supplemental Figure 4.** Heparin concentration was matched to physiologic concentration ($Hep_{High}$) and a tenth of the physiologic concentration ($Hep_{Low}$) by measuring the fluorescence of macrogels (n=3) on a Molecular Devices Confocal. Graph represents mean+/- standard deviation. Statistics: ANOVA. N=3.

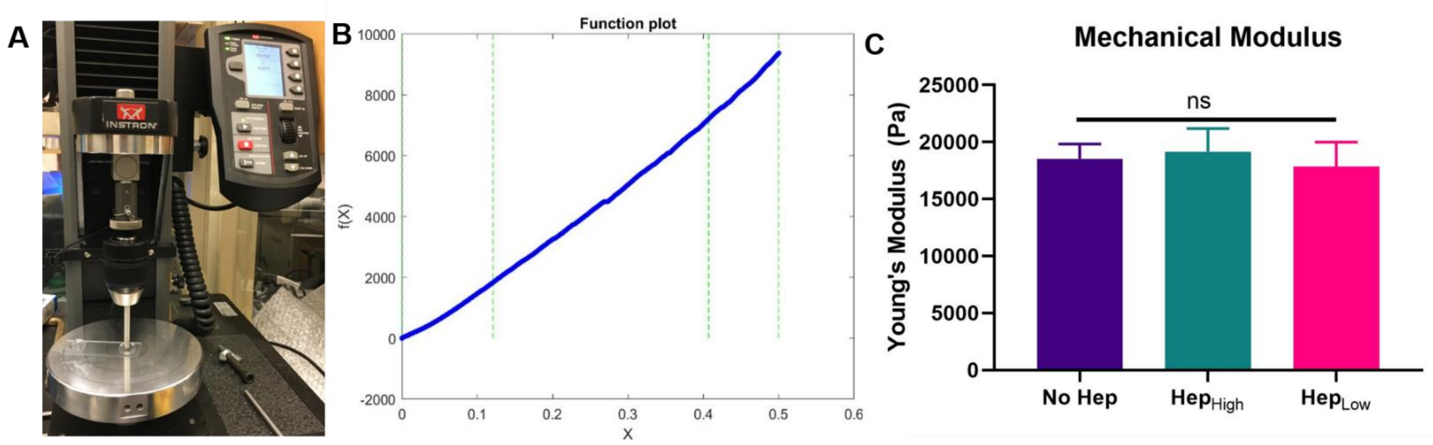


**Supplemental Figure 5:** Macrogels were **A)** tested on an Instron (n=3) and the **B)** Young's modulus is analyzed with MATLAB's shape language modeling package. C) Mechanical properties of the microgel types were matched to approximately 18kPa. Graphs represent mean+/- standard deviation. Statistics: ANOVA. N=3.

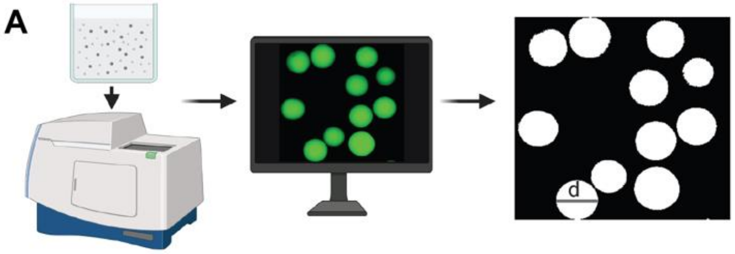


B

$$D_n = \frac{\sum N_i D_i}{\sum N_i}$$

$$D_w = \frac{\sum N_i D_i^2}{\sum N_i D_i}$$

$$PDI = \frac{D_w}{D_n}$$

C

| Particle Type | $D_n$ (µm) | $D_w$ (µm) | PDI |
|---|---|---|---|
| $Hep_{high}$ | 84.4 | 85.27 | 1.01 |
| $Hep_{low}$ | 90.59 | 93.10 | 1.03 |
| No Hep | 86.41 | 88.25 | 1.02 |

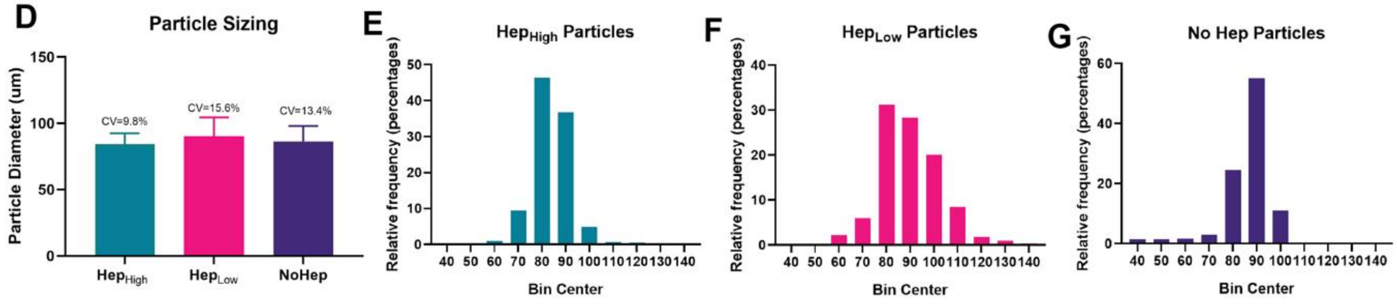


**Supplemental Figure 6: A)** Particles were generated using microfluidics and a dilute solution was imaged using a Molecular Devices Confocal. **B)** Diameters are calculated using a custom sizing module (ImageXpress) and **C)** PDI was calculated for each particle type. D) Average particle diameter for the three groups was between 84-90µm. Histograms of particle size distribution of **E)** $Hep_{High}$ particles, **F)** $Hep_{Low}$ particles, and **G)** No Hep particles.

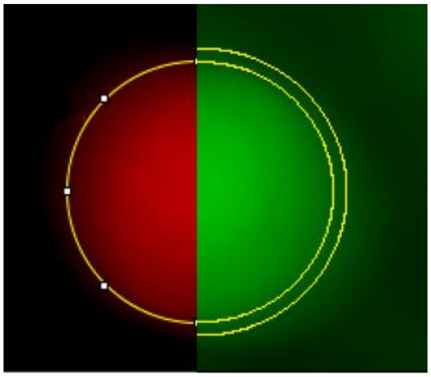

**Supplemental Figure 7:** To analyze gradient formation, particles are traced in their respective fluorescence channel and then that is overlayed on the streptavidin channel and the mean gray value is calculated in concentric circles surrounding the particle.

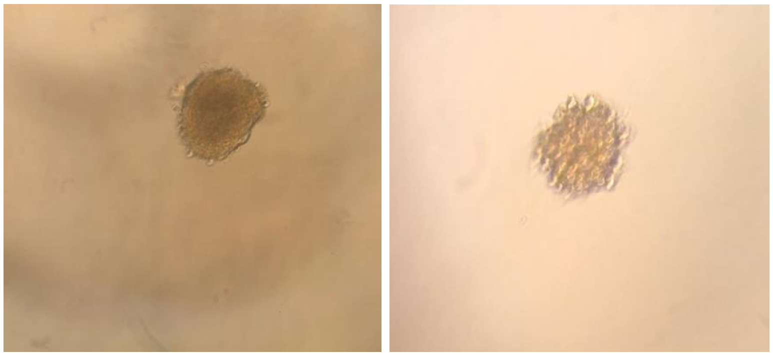

**Supplemental Figure 8***:* Human Dermal Fibroblast (HDF) and Human Dermal Microvascular Endothelial Cells (HDMVECs) spheroid formation after 24 hrs.

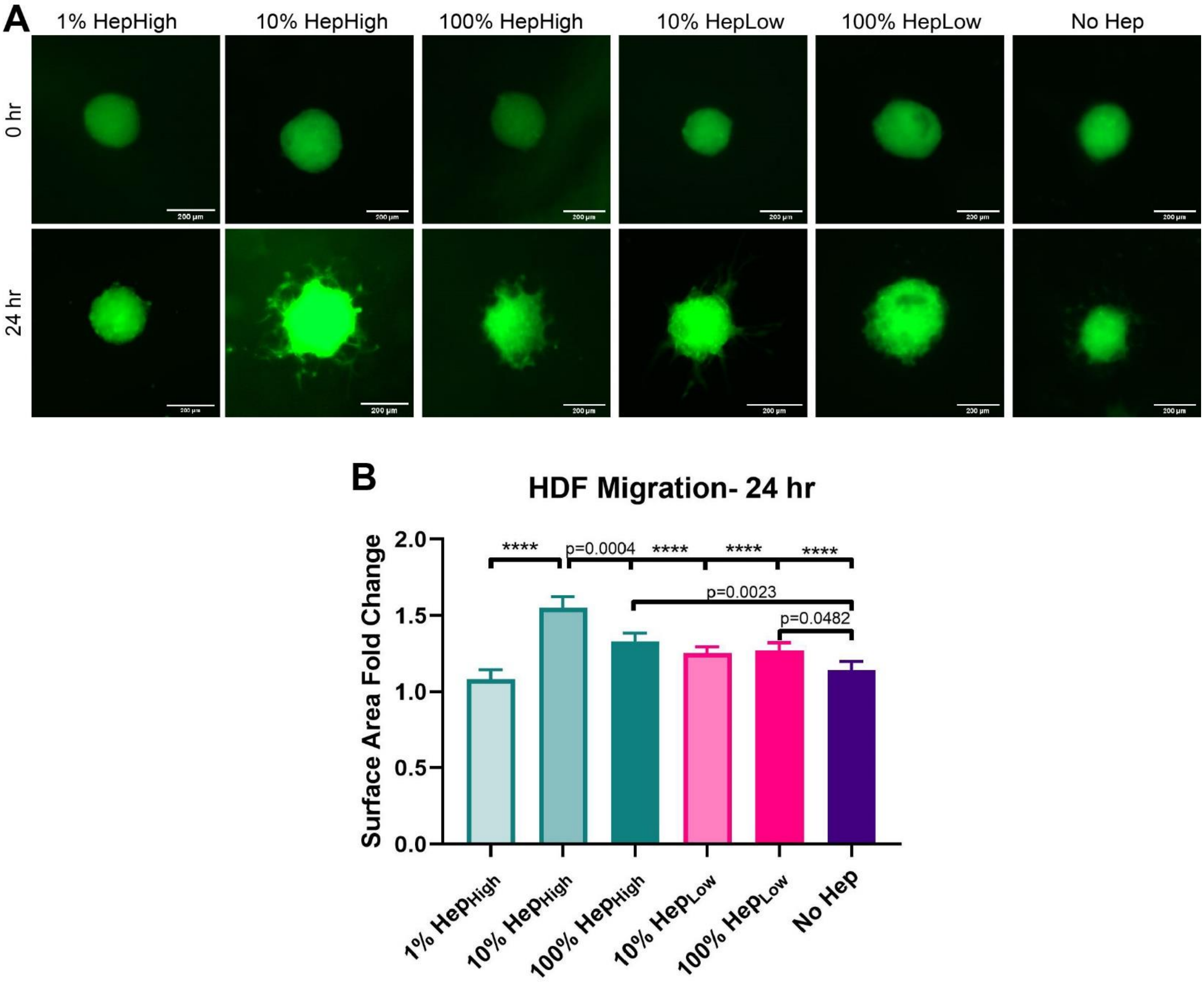


**Supplemental Figure 9: A)** Representative images of HDF spheroids at 0 hrs and 24hrs for each gel condition. **B)** Quantification of migration performed by analyzing the surface area fold change over 24 hrs. Graphs represent mean +/- standard deviation. Scale Bar represents 200µm. Statistics: ANOVA, Multiple comparisons post-hoc tests (Tukey HSD). N=4. Significance inside of bars represents comparison to no heparin gels. **** is p<0.0001.

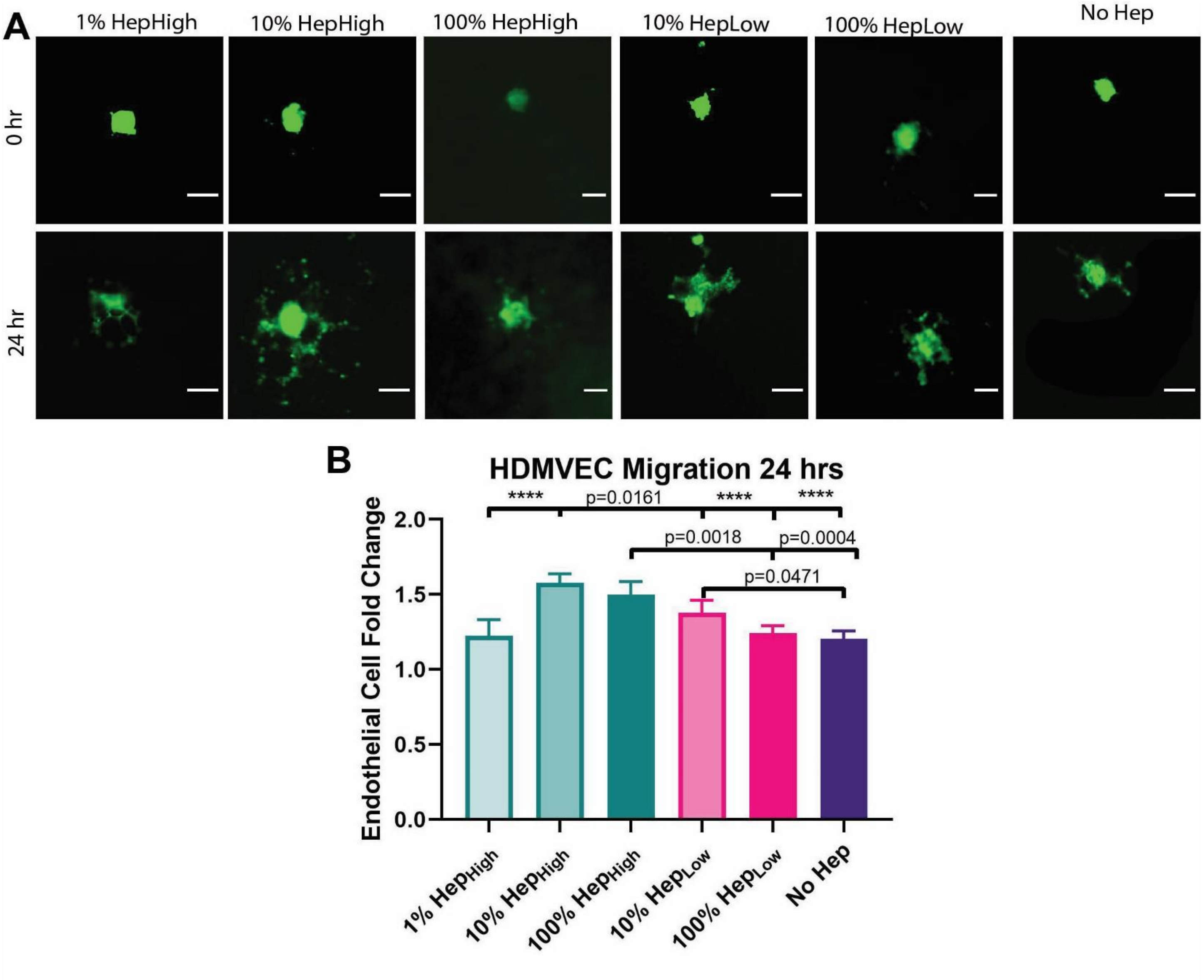


**Supplemental Figure 10: A)** Representative images of HDMVEC spheroids at 0hrs and 24 hrs for each gel condition. **B)** Quantification of migration performed by analyzing the surface area fold change over 24 hrs. Graphs represent mean +/- standard deviation. Scale Bar represents 50µm. Statistics: ANOVA, Multiple comparisons post-hoc tests (Tukey HSD). N=4. Significance inside of bars represents comparison to no heparin gels. **** is p<0.0001.

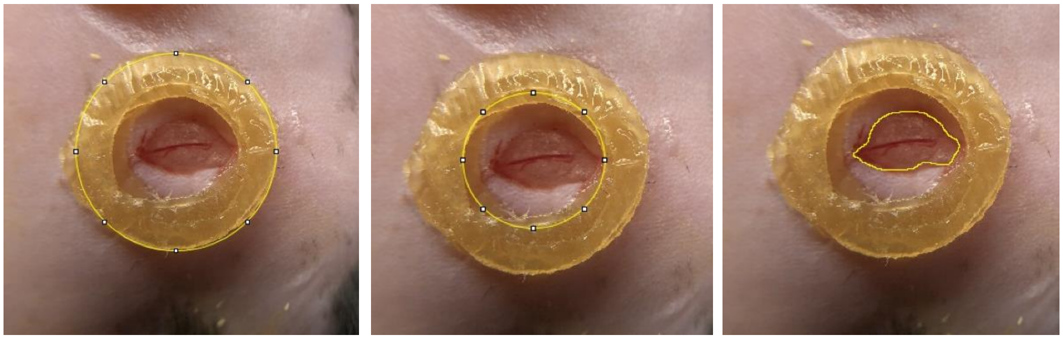

**Supplemental Figure 11***:* Image analysis to assess wound closure was accomplished using ImageJ. The area of the duoderm ring for the outer and inner edges was measured using the freehand trace tool.

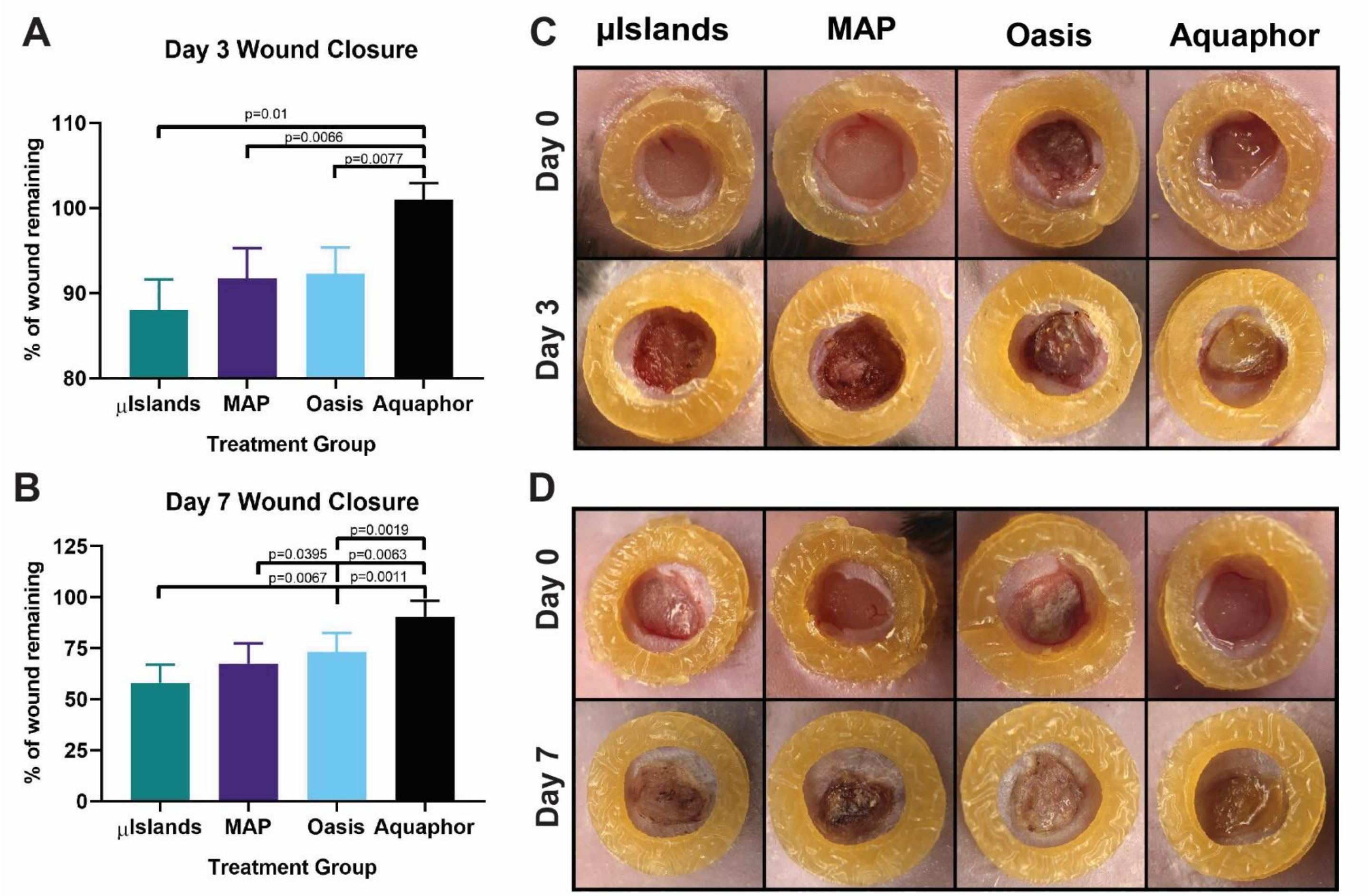


**Supplemental Figure 12:** Gross wound image closure at **A)** Day 3 and **B)** Day 7. **C)** Representative images of the splinted wounds at Day 0, 3 and 7 for each group. Graphs represent mean+/- standard deviation. Statistics: ANOVA, followed by a mixed model for multiple comparisons.

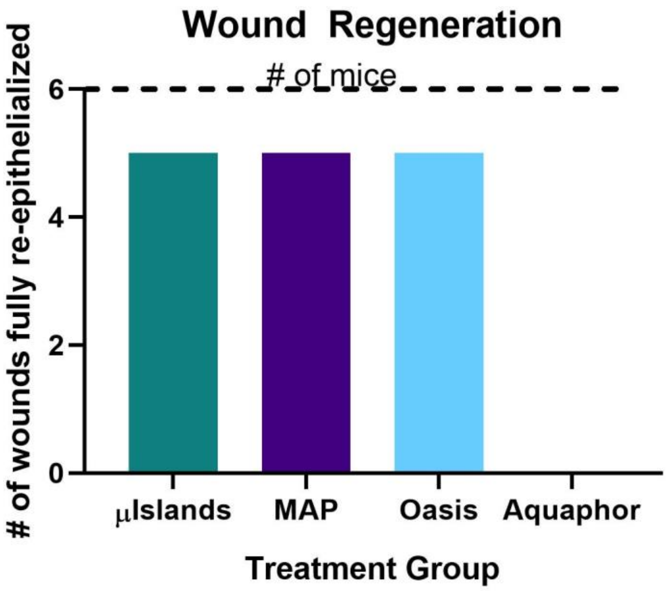


**Supplemental Figure 13:** Number of wounds fully re-epithelialized at Day 7 for each group.

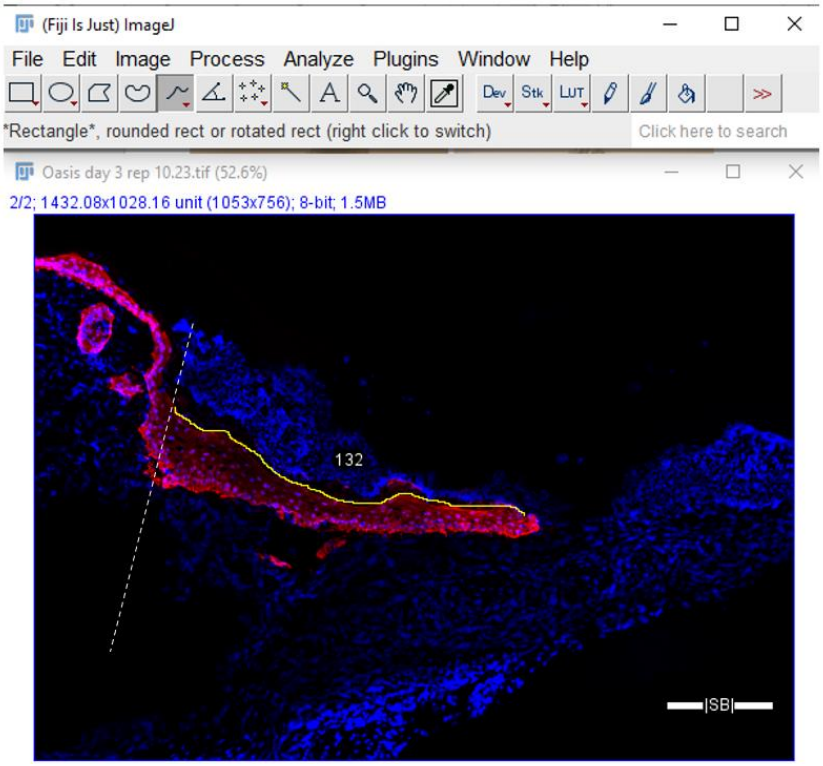


**Supplemental Figure 14:** Example analysis of epidermal tongue length. Epidermal tongue length was quantified by first setting the scale on ImageJ to convert measurements from pixels to microns. Next, the wound edges were marked by dotted white lines, as indicated by a change in tissue morphology from surrounding skin. Next, the freehand line tool was used in ImageJ to trace the top of the keratin-14 stain and the length was measured.

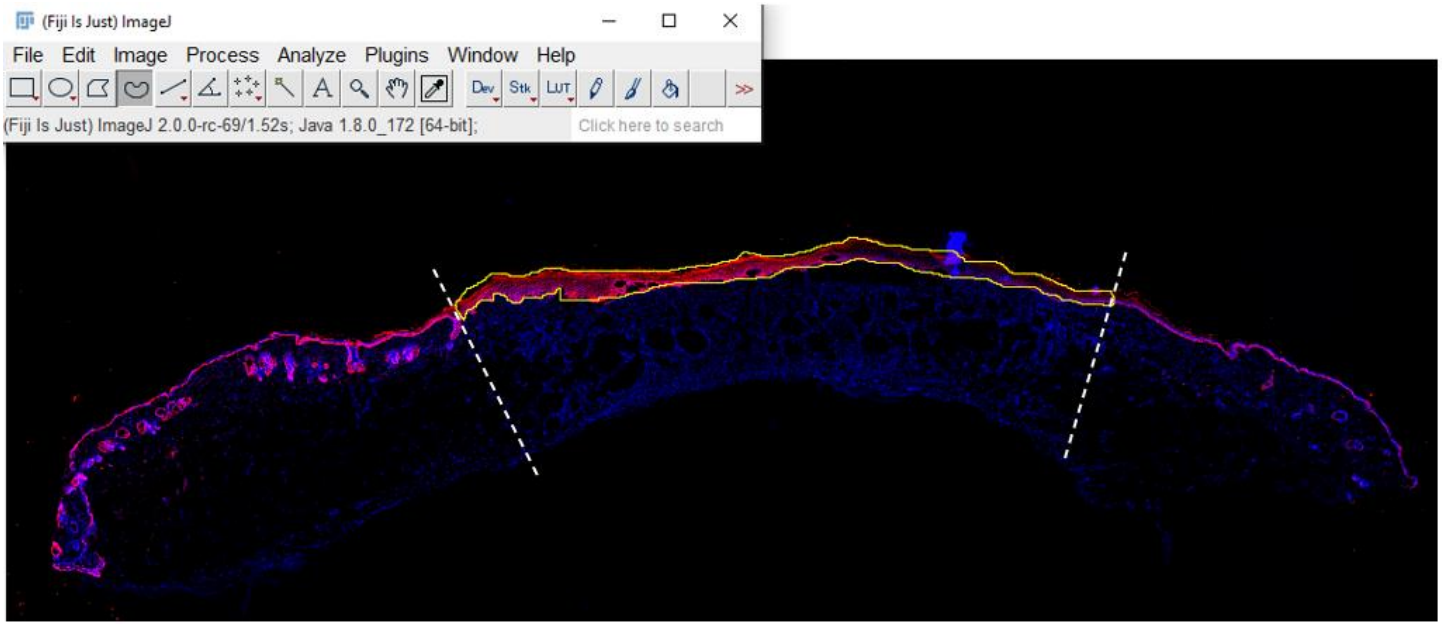


**Supplemental Figure 15**: Example analysis of epidermal thickness. Epidermal thickness was quantified by first setting the scale on ImageJ to convert measurements from pixels to microns. Next, the entire epidermal layer, as indicated by positive Keratin-14 staining (red) was outlined using the trace tool. Wound edges are marked by dotted white lines and were determined by changes in tissue morphology from surrounding skin to wound area. The area and width of the traced area were measured, and thickness was calculated by area divided by width.

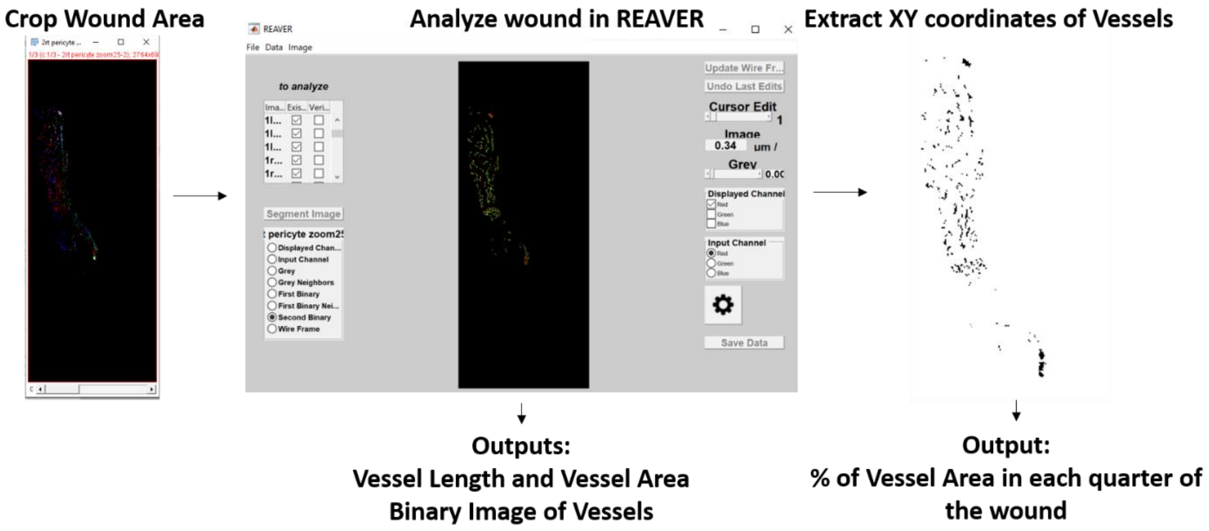


**Supplemental Figure 16**: REAVER analysis of blood vessels pipeline. Wounds were cropped in ImageJ to remove the surrounding skin, as determined by morphologic changes. The wound files were uploaded into the REAVER gui[32] on MATLAB and batch analysis was run. After analyzing, REAVER exports binary images for each image of the vessels and a table containing the vessel length and vessel area. The binary image's XY coordinates were exported. The wound length was calculated from the XY coordinates. Next, the wound length was divided into 4 sections, each representing 25% of the wound. From there the percentage of coordinates in each region was calculated to get the % vessel area in each quarter of the wound. This percentage was multiplied by the vessel length calculated from the REAVER to get the length of vessels at the edges or middle (inner 50% of the wound). These numbers were normalized to wound length for accurate comparisons between wounds.

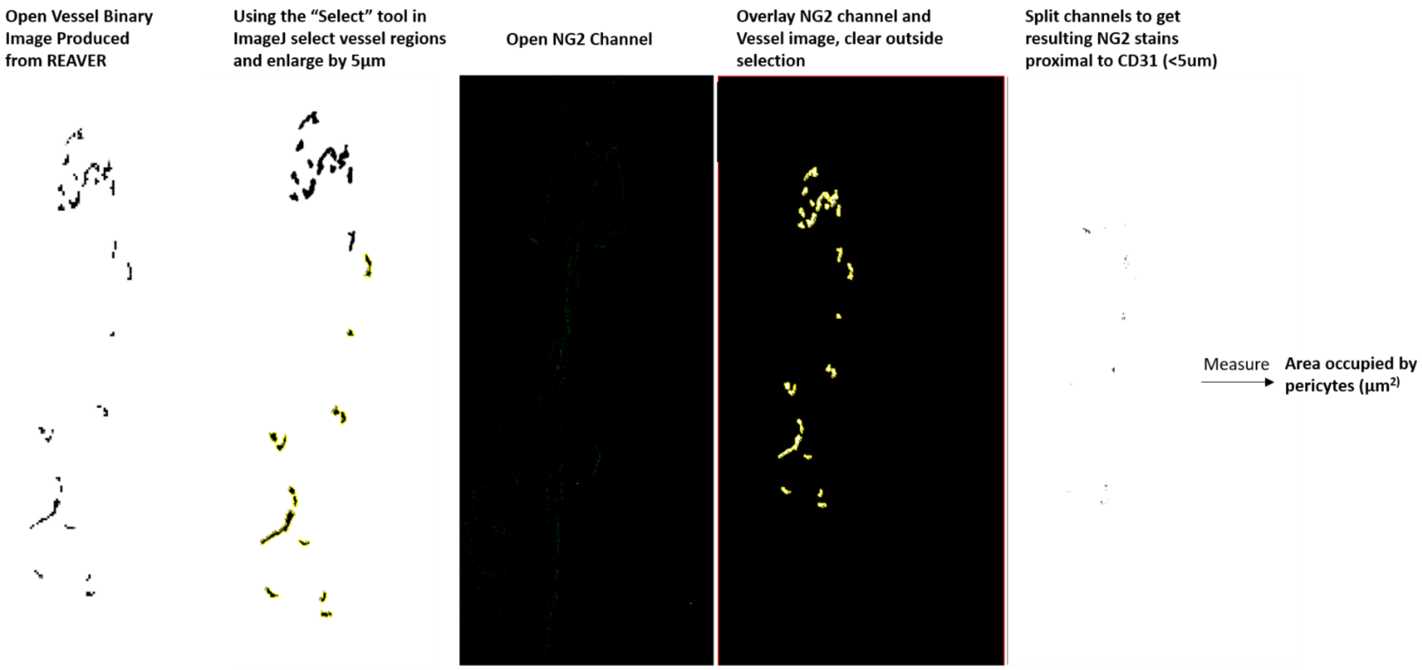


**Supplemental Figure 17:** Pericyte Analysis pipeline. Vessel binary images exported from REAVER were opened in ImageJ. Using the select tool, all vessels were selected and enlarged by 5µm to account for the area that pericytes could be occupying. Next, the NG2 channel image was opened and overlayed on the vessel image that had the pericyte regions. All staining outside of this region was cleared to remove any NG2 staining of other cell types. The channels were split to get the resulting NG2 channel that was positive pericyte staining. The area was measured to get the area occupied by pericytes in microns. This was normalized to the wound area in mm$^2$ which was previously measured by tracing on ImageJ to allow accurate comparison between wounds.

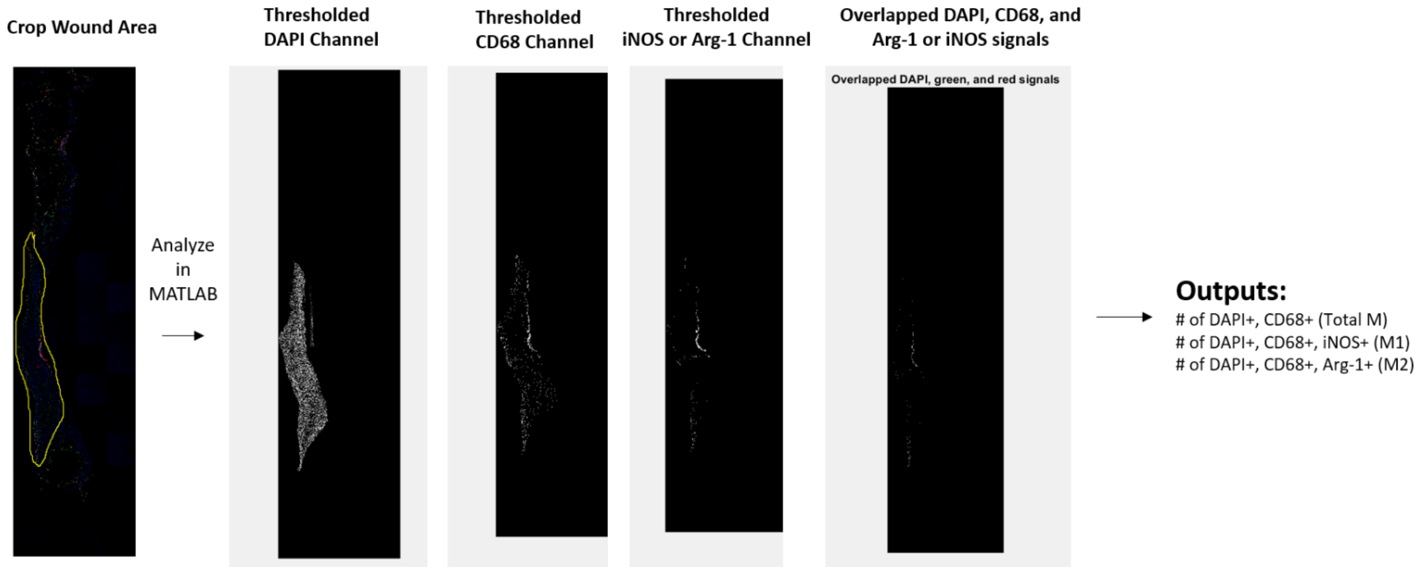


**Supplemental Figure 18:** Macrophage Analysis pipeline. Wound images from macrophage images were cropped using ImageJ to only have the wound area and not surrounding skin. Additionally, background removal was performed in ImageJ to remove any staining artifacts. This image was imported into MATLAB thresholded for each channel to determine the positive staining for DAPI, CD68 and either iNOS or Arg-1. The MATLAB code determined regions of overlap and total macrophages was calculated to be the number of cells with DAPI and CD68 co-localization. The amount of M1 macrophages was determined by the number of cells that had co-localization of DAPI, CD68 and iNOS. The amount of M2 macrophages was determined by the number of cells that had co-localization of DAPI, CD68 and Arg-1.

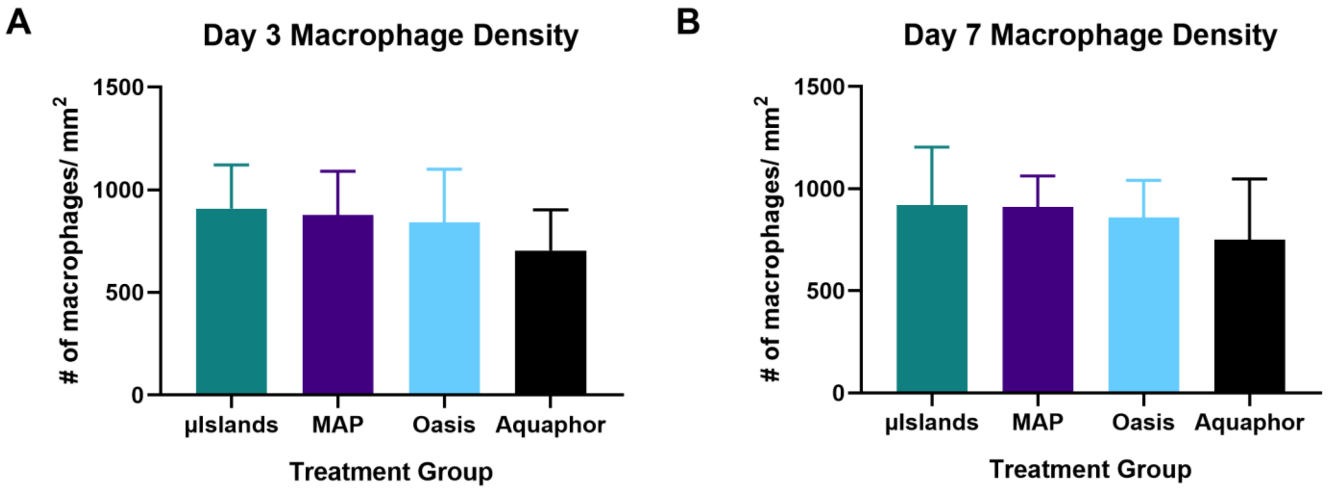


**Supplemental Figure 19**: Macrophage density quantified for each group at **A)** Day 3 and **B)** Day 7.

**Supplemental Table 1: Antibodies**

| **Antibody Name** | **Catalog #** | **Dilution** | **Secondary:** |
|---|---|---|---|
| Keratin-14 | BioLegend 905303 | 1:100, 5% goat serum | Goat Anti-Rabbit |
| CD31 | BioLegend 102516 | 1:50, 5% goat serum | None, conjugated with AF 647 |
| CD68 | BioLegend 137001 | 1:100, 5% goat serum | Goat Anti-Rat |
| iNOS | Abcam ab213987 | 1:100, 5% goat serum | Goat Anti-Rabbit |
| Arg- 1 | Santa Cruz Biotechnology SC-271430 | 1:50, 5% goat serum | Goat anti-Mouse |
| PDGFRB | Invitrogen MA5-15143 | 1:100, 5% goat serum | Goat anti-Rabbit |
| NG2 | R&D Systems MAB6689 | 1:50, 5% goat serum | Goat anti-Rat |
